# The metal cofactor zinc and interacting membranes modulate SOD1 conformation-aggregation landscape in an *in vitro* ALS Model

**DOI:** 10.1101/2020.07.25.220962

**Authors:** Achinta Sannigrahi, Sourav Chowdhury, Bidisha Das, Amrita Banerjee, Animesh Halder, Athi N. Naganathan, Sanat Karmakar, Krishnananda Chattopadhyay

## Abstract

Aggregation of Cu-Zn superoxide dismutase (SOD1) is implicated in the motor neuron disease, ALS. Although more than 140 disease mutations of SOD1 are available, their stability or aggregation behaviors in membrane environment are not correlated with disease pathophysiology. Here, we use multiple mutational variants of SOD1 to show that the absence of Zn, and not Cu, significantly impacts membrane attachment of SOD1 through two loop regions facilitating aggregation driven by lipid induced conformational changes. These loop regions influence both the primary (through Cu intake) and the gain of function (through aggregation) of SOD1 presumably through a shared conformational landscape. Combining experimental and theoretical frameworks using representative ALS disease mutants, we develop a ‘ **co-factor derived membrane association model**’ wherein mutational stress closer to the Zn (but not to the Cu) pocket is responsible for membrane association mediated toxic aggregation and survival time scale after ALS diagnosis.

## Introduction

The aggregation of SOD1 is believed to be one of the chief causative factors behind the lethal motor neuron disease Amyotrophic lateral sclerosis (ALS)^1^. Although more than 140 SOD1 mutations have been reportedly associated with ALS, there is no correlation between the stability (and aggregation) of these mutations and their disease manifestations. SOD1 aggregation has been investigated extensively *in vitro* by altering the solution conditions, such as temperature, pH, the presence of metal chelators, and the reduction of disulfide linkages^2, 3, 4^. The results of these studies clearly suggest that the aggregation of SOD1 is heterogeneous containing multiple steps, which is presumably the reason behind the lack of a structural understanding of aggregation processes^5, 6^.

WT SOD1 contains Cu^2+^ (Cu) and Zn^2+^ (Zn) as cofactors. It has been established that Cu is responsible for the primary function of SOD1 (the dismutase activity), and cell membrane acts as a scaffold in the process of Cu transfer to apo-SOD1 (metal free non-functional protein) through a Cu delivery chaperone (CCS)^7^. Previous studies have found noticeable presence of SOD1 in human serum lipoproteins, mainly in LDL and HDL, hinting at a possible protective role of SOD1 against the lipid peroxidation^8^. It has also been noted that SOD1 has a physiological propensity to accumulate near the membranes^9^ of different cellular compartments, including mitochondria, endoplasmic reticulum (ER) and Golgi apparatus^10^. In addition, computational studies have shown that the electrostatic loop (loop VII, residues 121–142) and Zn-binding loop (loop IV, residues 58–83) promote membrane interaction of apo-SOD1 initiating the aggregation process ^11^. Membrane binding induced aggregation of SOD1 has also been shown experimentally both in *vitro* and inside cells^6, 12, 13, 14^. Inclusions of SOD1 have been detected in the intermembrane space of mitochondria originating from the spinal cord^8^.

The above results can be reconciled by suggesting that cell membrane can play crucial roles not only in shaping up the primary function of the protein, but also in defining its aggregation process of generating fibrillar and non-fibrillar aggregates (the gain of function), with the loops IV and VII contributing critically to both processes. We hypothesize that (a) the induction of metal cofactors for the stabilization of loop IV and VII, membrane interaction and SOD1 aggregation would be some of the crucial elements in defining the overlapping folding-aggregation landscape of SOD1; (b) metal pocket perturbation by mutational stresses (as in disease variants) would modulate membrane association and facilitate aggregation, (c) the difference in aggregate morphology as a result of differential membrane interaction may contribute to the variation in cellular toxicity observed in ALS.

In this paper, we investigated the above hypothesis by studying how different structural elements (*i.e*. co-ordination of individual metals, membrane association and the location of mutations) attenuate the toxic gain of function of SOD1. An effective understanding of the role of individual metals (Cu and Zn) would require studying SOD1 variants containing only one metal (Cu or Zn) in addition to a variant that contains none. We have therefore prepared an apo (metal free) protein, which serves the latter purpose. For the former, we have generated two single metal containing mutants of SOD1, *viz*. H121F (only Zn, no Cu) and H72F (only Cu, no Zn), which are situated near the key loop VII (H121F) and loop IV (H72F) at the protein structure (Fig. 1a).

**Figure 1.**
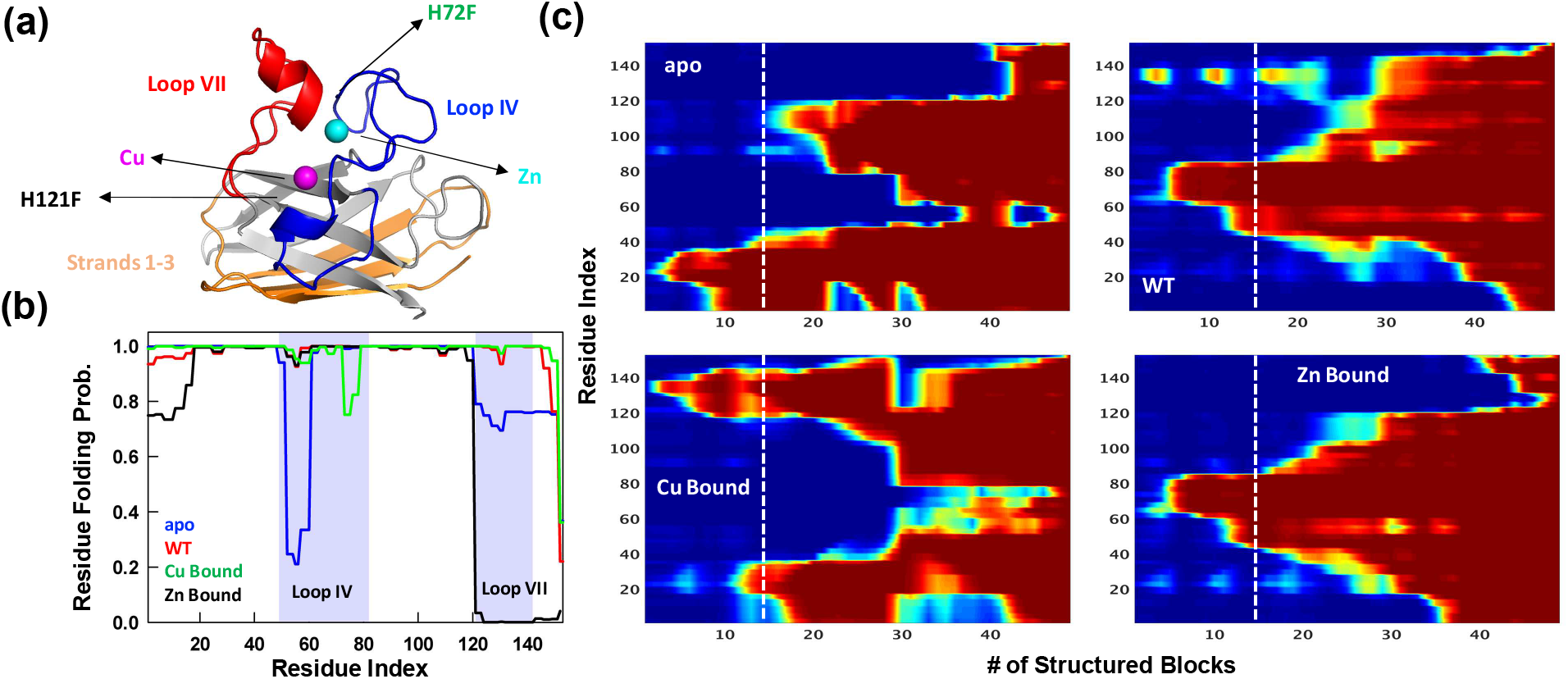
Statistical Mechanical Modeling of SOD1 Folding Mechanism. (a) Cartoon representation of SOD1 monomer highlighting the various structural elements. The positions of the mutation for H121F and H72F have been arrow marked. (b) Residue folding probability as a function of residue index for the different variants of SOD1 as predicted by the bWSME model. Note that WT represents the variant in which both the metallic cofactors are bound. (c) Average folding probabilities colored in the spectral scale going from 0 (dark blue) to 1 (dark red) as a function of the reaction coordinate, number of structured blocks. The vertical white dashed line signals the parts of the protein that fold first. For example, it can be seen that residues 1-40 fold early in the apo SOD1 (dark red) when compared to WT where residues 40-80 fold first.

Using computational analysis based on statistical mechanical model and detailed *in vitro* experiments, we propose here a “**Co-factor derived membrane association model”** of SOD1 aggregation and its possible implication in ALS. We demonstrate that differential metal binding and membrane assisted conformational changes can work in concert to attenuate the rate and propensity of aggregation. While apo (no metal) protein and H72F mutant (no Zn) experience strong membrane interaction, the WT (both metals) protein and H121F (no Cu) mutant do not show significant binding. We further find that membrane induced aggregates of H72F and apo protein showed significantly higher toxicity in terms of cell death and model membrane deformation when compared to WT and H121F mutant. We finally check the validity of this model to ALS using computational and experimental studies. For the computational validation, we show using fifteen ALS disease mutants, that the distance between the mutation site and Zn correlates well with the membrane binding energy and patient survival time after disease diagnosis, while Cu site does not seem to have any prominent role. For the experimental study we use two well studied disease mutants (G37R where mutational site is close to the Cu pocket and I113T where mutational site is near Zn pocket) to show that the model accounts well for their membrane binding/aggregation, correlating well with their disease onset phenotypes. This model puts forward a mechanism that Zn pocket destabilization (either by metal content variation or by mutational stress near Zn center) is the driving force behind the toxic gain of function of SOD1 mediated by the process of membrane association.

## Results

### Statistical Mechanical Modeling of SOD1 Folding Mechanism Hints at Aggregation Origins

The large size of SOD1 (151 residues) precludes a detailed characterization of the conformational landscape, the role of ions in determining the stability-folding mechanism and the effect of numerous mutations via all-atom simulation methods. To overcome this challenge and to obtain a simple physical picture of how the energetics of folding is governed by metal ions, we resort to constructing the folding landscape of reduced SOD1 variants through the statistical mechanical Wako-Saitô-Muñoz-Eaton (WSME) model (see Methods for model description and parametrization)^15, 16^. Here, we employ the bWSME model where stretches of three consecutive residues are considered as a block (b) that reduces the total number of microstates from 42.7 million to just ~450,000 ^17^.

The model, however, incorporates contributions from van der Waals interactions, simplified solvation, Debye-Hückel electrostatics, excess conformational entropy for disordered residues and restricted conformational freedom for proline residues^18, 19^. The predicted average folding path of SOD1 WT (with both Cu and Zn bound) highlights that the folding is initiated around the metal binding regions with early folding of the loop IV (nucleated by Zn) in the unfolded well and aided by flickering structure in the electrostatic loop (loop VII, nucleated by Cu). The rest of the structure coalesces around this initial folding site leading to the native state. This folding mechanism is very similar to that proposed earlier *via* detailed kinetic studies ^20^. In the absence of metal ions, the apo variant folds through an alternate pathway wherein the folding is nucleated through the first three strands (residues 1-40) following which the rest of the structure folds, recapitulating the results of single-molecule experiments (Fig. 1)^21^. It is important to note that the first three strands exhibit higher aggregation propensity as predicted from different computational servers (Fig. S1). Interestingly, the folding mechanism of the Zn bound SOD1 (with no Cu bound) is similar to that of the WT hinting that Zn coordination promotes proper folding. On the other hand, the folding mechanism of the Cu-bound SOD1 (in the absence of Zn) is similar to that of apo-SOD1 with additional folding probability in the region around the electrostatic loop. Taken together, the statistical modeling highlights how the absence of metals and particularly the absence of Zn (or mutations that affect Zn binding and not Cu binding) alters the folding mechanism by populating partially structured states involving beta strands in the unfolded well thus possibly increasing the chances of aggregation. Importantly, the model provides multiple testable predictions on the differential roles of Zn and Cu, which we address below *via* experiments.

### Cu deficient H121F behaves like WT SOD1 whereas Zn deficient H72F behaves like apo

We have recently shown that the mutants H121F and H72F contain negligible Cu and Zn respectively, while the apo protein is completely devoid of metal^22^. We validated this further using atomic absorption spectroscopy (Table S1) and activity measurements (Fig. S2a). We used steady state tryptophan fluorescence, far UV CD and FTIR spectroscopy to characterize these different proteins. SOD1 is a single tryptophan protein (Trp32), in which the tryptophan residue has been shown to be partially buried^23^. The role of Trp32 within the sequence segment ^30^KVWGSIKGL^38^ of high aggregation propensity has been investigated before^24^. We found that the formation of apo form resulted in a large shift in Trp32 emission maximum (332 nm for WT protein and 350 nm for apo protein) (Fig.2a). In contrast, other two mono-metallated variants (H121F and H72F) exhibited fluorescence emission maxima at wavelengths, which were intermediate between the WT and apo proteins (342 nm for H121F and 345 nm for H72F) (Fig.2a). Subsequently, we performed acrylamide quenching experiments to measure the solvent surface exposure of Trp32 for all variants. The values (Table S2) of the Stern-Volmer constant (*K_sv_*) were determined using a straight line fit, as shown in Fig. S2b. *K_sv_* for WT (6.8±0.1 M^−1^) was significantly lower than that of apo SOD1 (14.3±0.1 M^−1^). Both steady state fluorescence maxima and acrylamide quenching data clearly suggested an appreciable conformational alteration as the protein moved from its WT to the apo form. Interestingly, Zn-starved H72F mutant showed higher *K_sv_* compared to Cu-starved H121F mutant (Fig.S2b, Table S2).

Far UV Circular Dichroism (CD) data for WT and apo protein matched previous reports (Fig. S3)^25^. We used FT-IR spectroscopy to complement far UV CD results, using amide-I FTIR spectral region. The carbonyl (C=0) stretching vibrations at amide-I region provides information related to the secondary structure (beta sheet 1633-1638, alpha helix 1649-1656, disorder and turns and loops 1644 and 1665-1672 cm^−1^). Analysis of the secondary structure of WT protein (Fig. 2b) showed the presence of 10% alpha helix, 38% beta sheet, and 52% of turns and loops including disordered stretches. The percentage of the secondary structure determined from the FT-IR analysis was found to be consistent with the data obtained from the crystal structure (PDB 4BCY with 11% alpha helix, 40% beta sheet and 49% turns and loops) thus validating our method^26^. FT-IR data showed a decrease in beta sheet content (from 38% to 31%) as apo protein (Fig. 2c) formed. In contrast, the behavior of H72F mutant (Fig. S4a, beta sheet content of 32%) was found to be similar to the apo protein, while H121F mutant (Fig.S4b, beta sheet content of 37%) remained similar to WT protein. The percentage of secondary structure elements of all protein variants are shown in Fig. 2d.

**Figure 2:**
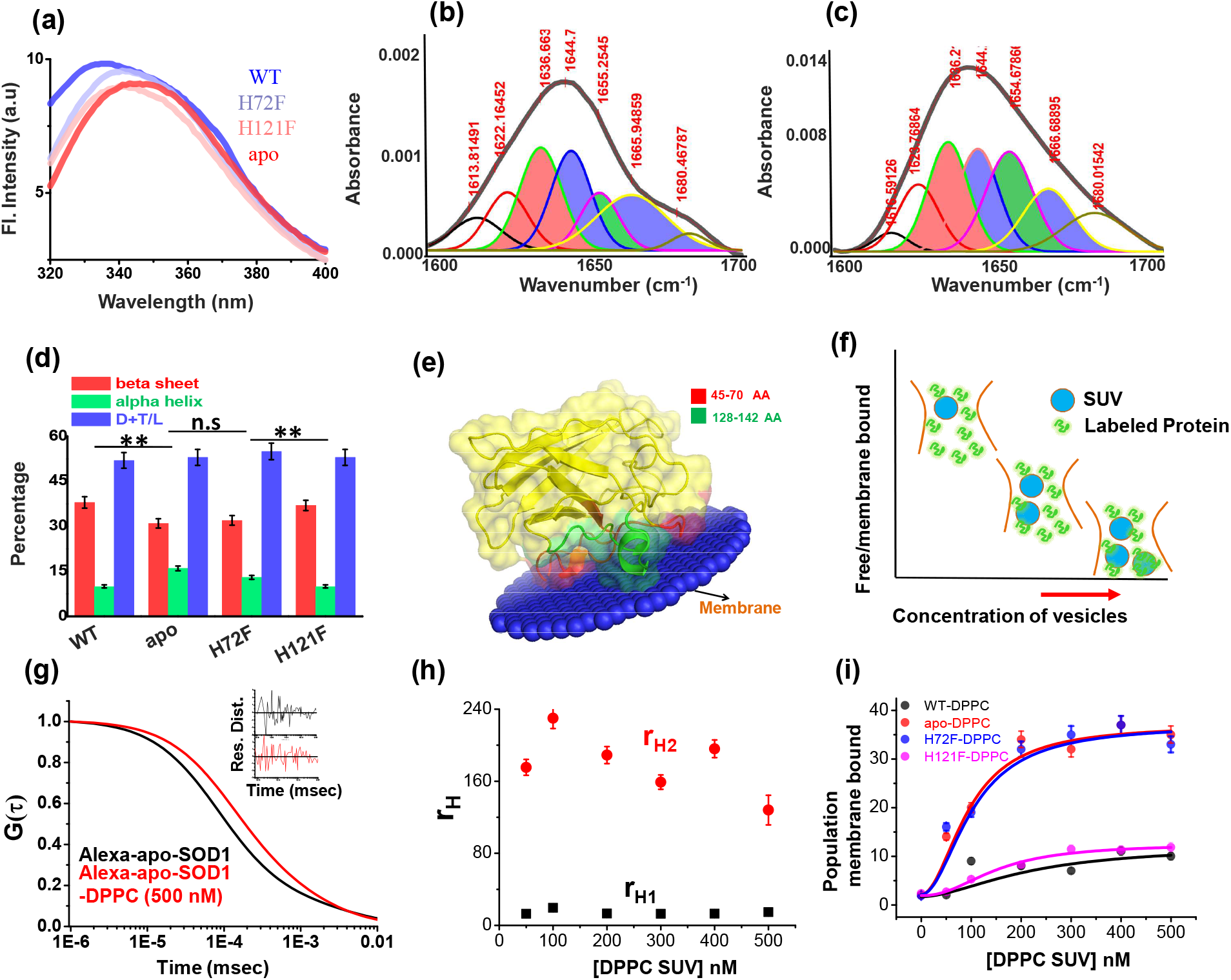
Structural characterization of SOD1 mutants and membrane association. (a) Steady state tryptophan fluorescence spectra of WT, apo and other two metal mutants (H121F and H72F). The WT displays an emission maximum at 332 nm, whereas the apo variant shows a red-shifted spectrum with the emission maximum at 350 nm. On the other hand, H121F and H72F show emission maxima at intermediate wavelengths. Deconvoluted FTIR spectral signatures of (b) WT and (c) apo. Red contour (~ 1637 cm^−1^) indicates beta sheet,blue color contour stands for disorder (1644 cm^−1^) and loops and turns (~1667 cm^−1^); green contour represents alpha helical character.All these secondary signatures were obtained by considering the amide-I spectra, which arises due to carbonyl frequency (C=O). (d) Percentage of different secondary structural components in WT, apo, H121F and H72F are shown in this figure. n.s denotes nonsignificant change while ** stands for significant change with P value<0.01. Error bars indicate the standard deviation of the data. Here, D+T/L stands for Disorder+Turns/Loops. (e) The membrane association of the apo protein as suggested by the OPM calculations. The membrane association of apo protein through the stretches 45-70 and 128-142 has been evaluated from the calculations. (f) A schematic representation of the membrane binding experiments using FCS, which suggests that with increasing concentration of DPPC SUVs, the population of the alexa labeled free monomeric protein (fast component) decreases with an increase in the membrane bound labeled protein (the slow component). (g) The correlation functions of alexa 488 maleimide labelled apo SOD1 in absence (black) and presence of DPPC SUVs (red). The inset shows the residual distributions of the correlation curves. (i) the hydrodynamic radii of monomeric apo SOD1 and membrane bound apo SOD1 were plotted against the concentrations of added DPPC SUVs. The average hydrodynamic radius of fast component i.e free monomeric apo SOD1 (r_H1_) was found to be 13.5 Å whereas the average radius for slow membrane bound protein molecule (r_H2_) was found to be 170 Å. We did not see any change in the values of r_H1_ and r_H2_ with increasing DPPC SUV concentrations. (i) amplitude percentages of the membrane bound protein variants were plotted against the concentrations of DPPC SUVs. The solid lines through the data were used to determine the values of K_a_ for each protein variant.

### Zn deficient SOD1 shows higher membrane association compared to the Cu deficient and WT proteins

To obtain a preliminary understanding of the possible membrane binding sites of SOD1, we resorted to computational techniques using ‘Orientation of protein in membrane’-tool^27^, which predicted weak interaction of WT on membrane surface (Fig. S5a). In contrast, the same calculation predicted higher binding affinity of apo protein with the membrane (Fig.2e). The computed values of ΔG_transfer_ (free energy change of protein transfer from bulk to the membrane) was found to be substantially higher for the apo (−2.6 Kcal mol^−1^) while compared to WT protein (−0.9 Kcal mol^−1^). We validated the above computational prediction using experimentally determined protein-lipid binding constants (K_a_, M^−1^), which we measured using fluorescence correlation spectroscopy (FCS). FCS monitors diffusional and conformational dynamics of fluorescently labeled biomolecules at single molecule resolution. ^28^ For FCS experiments, we labeled the cysteine residues of all the SOD1 variants using Alexa-488-maleimide. Fig. 2f shows a schematic diagram of how the labeled proteins and protein-lipids complex would behave inside the confocal volume. Using FCS we determined the correlation functions using 50 nM Alexa488Maleimide protein in the presence of increasing concentration of DPPC SUVs. We fit the correlation functions using a two component diffusion model and the goodness of the fit was established using the randomness of the residual distribution. Fig. 2g showed the typical correlation functions of alexa labeled apo SOD1in absence and presence of 500nM DPPC SUVs). In this model, the fast and slow diffusing components corresponded to the free (with r_H1_ 13.5Å) and lipid bound protein (r_H2_ 170Å) respectively (Fig. 2h). With increasing DPPC SUV concentration, the percentage of slow component increased (Fig. 2f), which occurred at the expense of the fast component, and a sigmoidal fit of either of these components yield the values of K_a_, which showed that the binding affinities followed the trend: apo ≥ H72F > H121F >WT (Fig.2i, Fig.S5b,Table S3).We then measured the Stern Volmer constants using acrylamide quenching experiments of Trp32 fluorescence with protein variants in the absence (K_sv_) and presence of (K_svm_) membrane. The parameter K_sv_/ K_svm_ was found maximum for the apo protein, and minimum for WT (Fig.S5c, Table S2). H121F and H72F variants behaved like WT and apo protein respectively. As observed by FT-IR, DPPC binding resulted in no or minimum change in conformation for WT and H121F proteins (Fig.S5d,e,h), while a large decrease in beta sheet content with simultaneous rise in non-beta content, specifically alpha helical content, was observed for the apo protein and H72F mutant (Fig.S5f,g,h).

### Lipid vesicles accelerate aggregation kinetics of apo and Zn deficient mutants

Aggregation kinetics of WT, apo and the mutant SOD1 in their TCEP reduced states were studied systematically both in the absence and presence of DPPC. A typical protein membrane ratio of 1:2 was maintained for all measurements involving membranes. For the initial assessment of the aggregation kinetics, the fluorescence intensity enhancement of amyloid marker Thioflavin T (ThT) was monitored. ThT is known to bind to protein aggregates with cross beta structure giving rise to a large increase in its fluorescence intensity. From the ThT fluorescence assay, we found that the WT protein does not aggregate, both in the absence or presence of membrane (Fig.3a,b). For the H121F variant in the absence of membrane, we found a small and slow enhancement of ThT fluorescence and the profile remained unchanged when we added the membrane (Fig. 3a,c). In contrast, for apo and H72F variants, ThT assay showed large fluorescence increase and the kinetics followed typical sigmoidal patterns. The addition of membrane increased the rate of aggregation for both variants and a large decrease in the lag times. When compared between the apo and H72F variants, we found that the rate of aggregation is higher (i.e. with less lag time) for the apo protein (Fig.3a,b,c Table S4).

**Figure 3:**
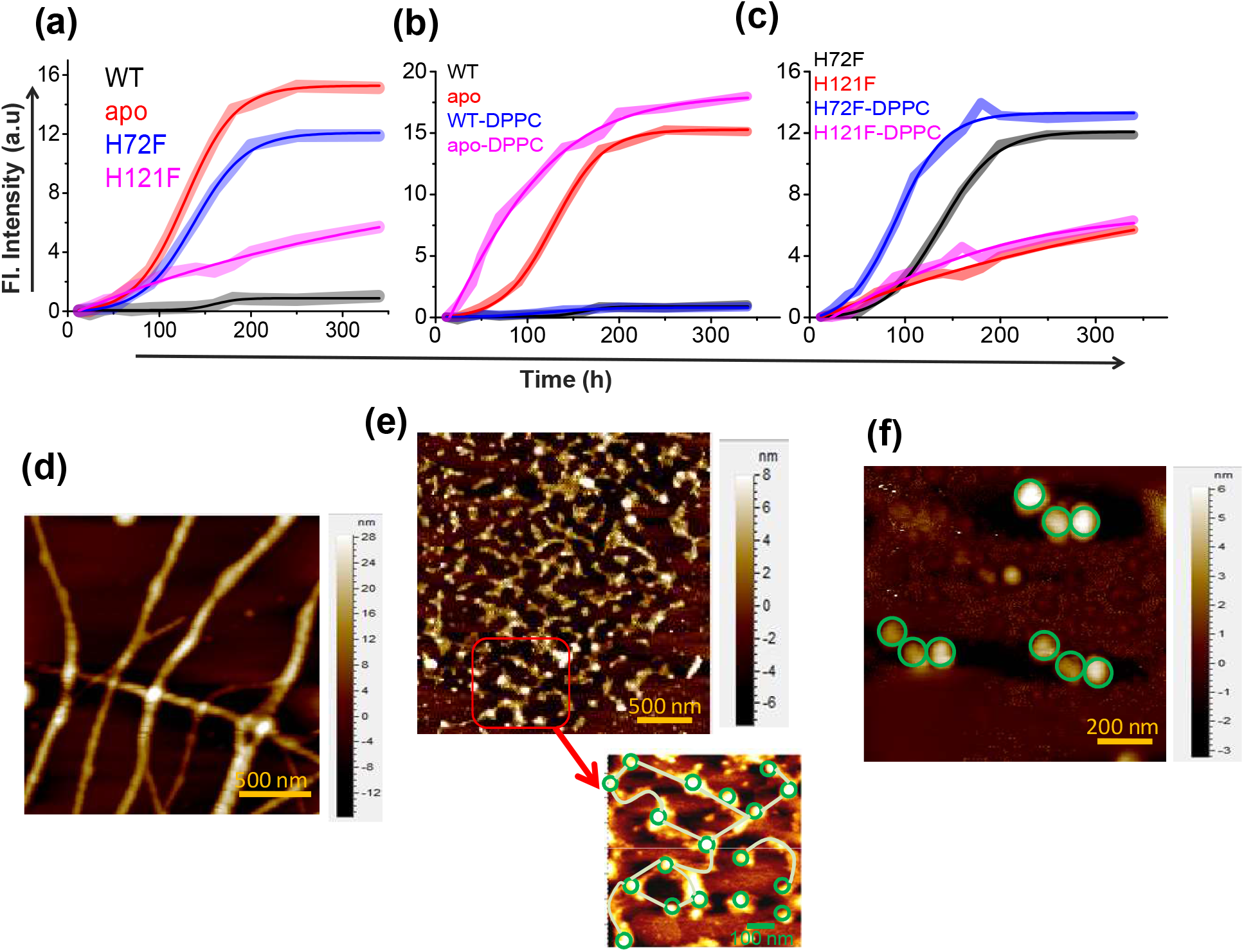
Aggregation of WT SOD1 and its mutants and membrane effects. (a) ThT fluorescence (at 484 nm) for the protein variants under reducing conditions to monitor the kinetics of aggregation. (b) Time-dependent increase in ThT fluorescence intensity of WT and apo both in the absence and presence of membrane (DPPC SUVs were used here as membrane). (c) Same as panel b but for H72F and H121F. AFM images of the aggregates of apo SOD1 in absence (d) and presence (e) of DPPC SUVs. These AFM images were taken at the plateau of the ThT aggregation curves. AFM images of apo_agg_ showed linear fibrillar aggregates with an average size 1.8-2 μm. In contrast in the presence of membrane (apo_aggm_), we found network of small fibrils, which were connected by DPPC vesicles (as drawn in the inset of Figure 3e). It may be noted that the size and height (average diameter is 70 nm and average height is 7 nm) of the connecting spherical objects are similar to (f) AFM micrograph of the control DPPC SUVs which showed distinct membrane structures with an average size of 70-90 nm.

We then imaged using atomic force microscopy (AFM) the aggregates collected from the plateau regions of the aggregation kinetics (at a time point when the fluorescence of THT was maximum (saturated) and did not change). Protein (P) aggregates will be denoted by P_agg_, and P_aggm_ to indicate if they are formed in the absence or presence of membranes respectively. For example, the aggregates of WT in the absence and presence of membranes would be denoted by WT_agg_, and WT_aggm_ respectively. AFM imaging also showed that in the absence of membrane, WT and H121F did not form aggregates, fibrillar or otherwise (Fig.S6), while large fibrillar aggregates were found to form with apo (Fig. 3d) and H72F mutant (Fig. S6). The average size of the fibrillar apo_agg_, was found to be 1.8-2 μm with an average height of 20 nm. H72F_agg_, showed similar morphology (Fig. S6). Significant morphological differences were noticed for the aggregates of apo and H72F variants, in the absence (Fig. 3d,e and Fig. S6) and the presence of the membrane. The apo_aggm_ appeared to exhibit network of thin aggregates (the average size was found to be 700-800 nm with an average height of 6-8 nm) which were found to be connected by the spherical DPPC vesicles (Fig.3e, inset; Fig. 3f).

### Aggregates of apo and H72F variants that form in presence of membrane show highest cellular toxicity and GUV deformation rates

We investigated the effect of the aggregates of different protein variants on cellular toxicity (by measuring cell viability) in general and on the cell membranes (using a number of spectroscopy and imaging assays) in particular. Cell viability was measured by the standard MTT assay. We used aggregates collected at the plateau region of the aggregation kinetics. MTT assay using SHSY5Y cell line showed minimum toxicity in terms of cell death for WT_agg_, WT_aggm_, H121F_agg_ and H121F_aggm_ (Fig. S7). In contrast, apo_agg_, and H72F_agg_, showed significantly higher neuronal dead cell population which increased further with apo_aggm_ and H72F_aggm_ respectively. Although neuronal cell death is one of the decisive factors of aggregate toxicity, the severity of ALS has been found to depend on the extent of membrane perturbations, which may contribute to multiple events including (i) mitochondria associated membrane (MAM) collapse and disruption ^29^ and (ii) the synaptic dysfunction due to impaired synaptic vesicles function towards neurotransmission^30, 31^ and (iii) the prion like spread of toxic aggregates between cells presumably through macropinocytosis^32, 33^. Therefore, it is necessarily important to investigate in detail the protein aggregates induced membrane-perturbation which has never been addressed before. We used three different assays for the *in vitro* studies, *viz*. a) phase contrast microscopy using a membrane model of Giant Unilamellar Vesicle (GUV) to probe how the presence of aggregates change their size and shapes, b) a calcein release assay to probe aggregate-induced pore formation, and c) FTIR to determine the molecular mechanism of the influence of different aggregates on the structure of lipids.

For the imaging assay, we used time based optical microscopic investigation to unveil how the addition of P_agg_, and P_aggm_ affects the size and shape of GUVs. GUV is widely used as a model membrane system, providing free-standing bilayers unaffected by support-induced artifacts yet with sufficiently low curvature to well mimic cellular membranes and mitochondrial membrane as well. An advantage of this assay comes from the use of phase contrast without requiring any external fluorophore label. We made the GUVs composed of DOPC:DOPE:PI:DOPS:CL in the ratio 4.5:2.5:1:0.5:1.5, which mimics mitochondrial membrane composition^34^. Fig. 4a is a representative example of how apo-aggregates formed in the absence of membrane (apo_agg_,) behaved with the GUVs. In contrast, Fig. 4b shows the influence of the apo-aggregates formed in the presence of membrane (apo_aggm_) (Fig. S8). Fig. 4c shows an example of a control, in which WT samples (WT_agg_, which should contain minimum aggregates population) were added to the GUVs. GUV images of Fig. 4a and 4b clearly demonstrate that the aggregates attach on the surface of GUVs (shown by arrows) leading to membrane-deformation and change in lamellarity. These two images also show the pore formation, which was further established by the contrast loss. A comparison between Fig. 4a and 4b shows visually that membrane deformation is more prominent (more damage) and faster (occurs at earlier time points) in the presence of P_aggm_. All these changes in GUVs were found absent in Fig. 4c, in which WT_agg_, was used. For the quantification of the image data, we determined the difference in refractive indices between the exterior and interior of GUVs (represented by I_ptp_, peak to peak intensity in Fig. 4d)^35^. In the case of intact vesicles, the values of I_ptp_ would be high (Fig. 4d, left) due to the difference in sugar asymmetry between outside and inside of GUVs, which would disappear in the case of membrane deformation (Fig. 4d, right).We plotted the time dependence of I_ptp_ to determine quantitatively the membrane deformation kinetics by measuring the deformation rate constants, λ (Fig. 4e, inset). The GUV deformation was insignificant and no detectable kinetics were found for WT_agg_, WT _aggm_, H121F_agg_, and H121F_aggm_ (Fig. 4e). On the other hand, apo_agg_. and H72F_agg_, exhibited significantly high deformation rate (λapo_agg_ ~1.8×10^−3^ s^−1^and λH72F_agg_~2.1×10^−3^ s^−1^) and both kinetics appeared cooperative (sigmoidal behavior, Fig. 4e). It is interesting to note that in presence of apo_aggm_ and H72F_aggm_ the deformation rate increased significantly (λapo_aggm_ ~3.5×10^−3^ s^−1^and λH72F_aggm_ ~3.9×10^−3^ s^−1^). More interestingly, both apo_agg_ and apo_aggm_ showed a tendency to create attachments between vesicles to generate vesicular assembly and co-operative deformations (high contrast images in Fig. 4f, which is also shown by a schematic drawing). The image analysis and visualization of the GUVs in presence of apo_aggm_ suggested that the co-operative vesicular clustering and deformation presumably occurred through allosteric communications mechanism by the aggregates (Fig. S8). We found that apo_aggm_ and H72F_aggm_ are more efficient towards inducing vesicular assembly and deformations (Fig. 4f, Fig. S9, S10).

**Figure 4:**
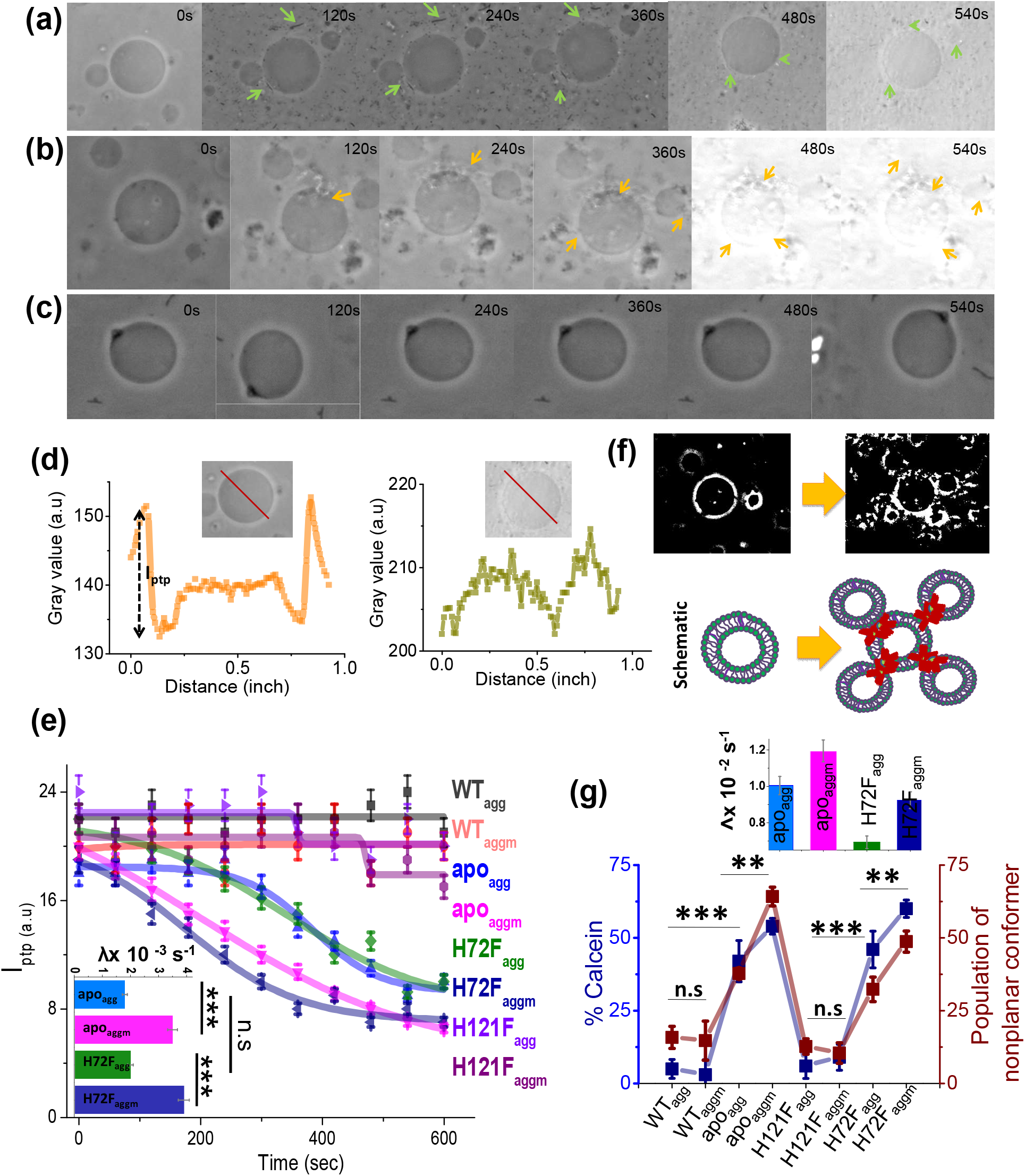
Membrane deformation by protein aggregates. Time variations of phase contrast micrographs of a single GUV when GUVs were treated with (a) apo_agg_, (b) apo_aggm_ and (c) WT_agg_. The images show gradual contrast loss, the loss of lamellarity and aggregate association with the vesicles for (a) and (b), while (c) does not show any change. The figure (b) also shows few vesicles assemblies as induced by aggregates (also refer to a high contrast image below). The protein aggregates and their association with GUVs have been marked by green and yellow arrows. (d) The pictorial definition of I_ptp_ and how I_ptp_ changes for an intact (left) and porus (right) vesicle. (e) The values of I_ptp_ are plotted for different protein aggregates with time. The inset of this figure shows the rate of deformation (λ, sec^−1^) for apo_agg_/ apo_aggm_ and H72F_agg_/ H72F_aggm_. The typical sizes of the GUVs were ~30 μm. (f) High contrast images of GUVs show vesicular assembly in presence of apo_aggm_, which is also schematically described in the figure below using a drawing. The schematic representation shows that apo_aggm_ can act as a connector between multiple GUVs. (g) Plot of calcein leakage percentage and the population of nonplanar rotamers (these rotamers arise at 1367 cm^−1^ vibrational frequency region of hydrocarbon lipid chains on treatment with different protein aggregates). Here, the subscripts agg and aggm stand for the aggregates of the respective protein species at the plateau region of the aggregation profiles which were formed in absence and presence of membrane (DPPC SUVs) respectively. The inset of this figure shows the rate of fluorescence growth(∧,sec^−1^) due to aggregate induced pore formation mediated calcein dye leakage from the SUVs that mimic synaptic vesicles (composition of lipid: DOPC:DOPE:DOPS in the ratio 2:5:3). Here n.s designates nonsignificant change whereas ** stands for significant (p value < 0.01) and *** for highly significant (p value < 0.001). The error bars indicate the standard deviation of triplicate experimental data.

Electron microscopic images of the synapses infused with WT SOD1 and G85R-SOD1clearly showed that G85R-SOD1-infused synapses showed vacant AZs (active zones) and occasional abnormal membranous structures whereas there occurred no reduction in the synaptic vesicles number in case of WT SOD1^31^. Similar observation was found previously by Wang etal^36^. Using C.elegans as model system, they showed that the neuronal toxicity in ALS appears due to synaptic dysfunction that occurs because of misfolded and aggregated disease mutant driven lowering in the number of organelles including synaptic vesicles and mitochondria. The pore formation and vesicle rupture by aggregates may be possible reason for the reduced synaptic vesicle population in case of ALS disease mutant. To investigate this issue, we prepared calcein entrapped SUVs composed of DOPE, DOPS and DOPC at the molar ratio 5:3:2 to mimic the synaptic vesicle composition and curvature^37^. In this assay, we measured the percentage of calcein leakage from the dye entrapped inside lipid vesicles (Fig. 4g, Fig. S11a)^35^.The extent of calcein leakage after the treatment of different protein aggregates followed the trend similar to what was observed by GUV micrographic observation (Fig. 4e) which suggested that the membrane rupture is facilitated in significantly higher rate and extent by apo_aggm_ and H72F_aggm_ while compared with apo_agg_ and H72F_agg_ respectively(Fig.4g, inset, Fig. S11a).

In order to induce membrane deformation as observed by previous two assays, the aggregates need to inflict substantial changes in lipid structure. To determine the extent of conformational change the lipid molecules experience by protein variants, we used ATR-FTIR to measure quantitatively the populations of different rotamers in a general planar trans-oriented phospholipid bilayer of DPPC. CH_2_ wagging band frequency (1280-1460 cm^−1^) of the hydrocarbon tail region of the bilayer was carefully monitored for this purpose.^38, 39^ The results show a significant increase in the populations of nonplanar kink+gtg^/^ rotamers (this band appears at 1367 cm^−1^) when planar lipid bilayer was treated with apo_agg_ /apo_aggm_ and H72F_agg_/H72F_aggm_ (Fig. 4g; Fig. S11b-c). We also found that apo_aggm_/H72F_aggm_ exerted greater effect on lipids than apo_agg_,/H72F_agg_,. Interestingly, differences in the populations of non-planar conformers are found to be similar to the variation in the extent of calcein release induced by protein variants (Fig. 4g). This observation provides preliminary evidence that both events occur by similar trigger (presumably the membrane attachment by the aggregates).

### Aggregation kinetics and aggregate induced toxicity studies of ALS disease mutants

In the previous few sections using WT, apo, H121F and H72F we established that the WT (containing both Zn and Cu) and the Zn bound H121F protein did not aggregate or induced toxicity through membrane deformation. In contrast, the removal of either Zn (H72F mutant) or both metals (the apo protein) resulted in strong membrane association, aggregation and induction of cellular toxicity. The data clearly suggest that Cu plays secondary roles in aggregation induced toxicity, while Zn pocket destabilization acts in concert with membrane-induced conformational change resulting in aggregation and toxic gain of function. Since these experiments validate successfully the predictions from the statistical mechanical model, we subsequently wanted to understand this behavior could be generalized in ALS disease mutants. For this purpose, we used both *in silico* and experimental approaches. For the *in silico* method, we selected fifteen disease mutants of ALS, whose structures are available in the protein data bank (PDB) (Table S5). From the crystal structures, we determined the distance between the mutation site and Zn (and Cu) for all these disease mutants. In addition, we calculated the transfer free energies, *i.e*. the theoretical membrane association energies (ΔG_Tr_), of these mutants using the OPM server. We found that the negative values of ΔG_Tr_ decreased linearly with the increased distance of Zn site for these mutants, while it remained non-variant with the distance of Cu site (Fig. 5a). We also found that the severity of ALS mutants (defined here as the survival time in year of ALS patients after diagnosis) correlates nicely with the distance of the Zn sites (Fig. 5b)^40^. For the experimental validation, we used two representative ALS disease mutants of varying distance between the mutation site and Zn. The mutations, namely G37R (mutation site-Zn distance 24Å, less severe) and I113T (mutation site-Zn distance 16Å, more severe) (Fig. 5b) are well studied in literature^41, 42, 43^. Using FCS we also measured the K_a_ values of these two disease mutants which showed greater binding affinity for I113T (Fig.5c, Table S3). Structural characterization using FTIR showed greater extent of alpha helical content in I113T compared to G37R (Fig. S12). On the other hand, further increase in the alpha helical structure in I113T was observed on interaction with model membrane. In contrast, we did not observe any membrane induced structural change for G37R (Fig. S12a-e). To understand the internal structural changes due to the disease mutations, we performed some *in silico* study using DynaMut webserver (http://biosig.unimelb.edu.au/dynamut/) to predict the effects of mutational stress on the conformational dynamics, stability and flexibility of protein.^44^ The results suggested that G37R mutation increased the rigidity of different regions (6-16, 30-42, 80-86 aa) of the protein whereas significant increase in flexibility of the loop IV and VII region (48-52,130-150) was observed in case of I113T mutation (Fig. S13). The predicted changes in folding free energy (*ΔΔG, kcal/mol*) for I113T was found to be much more negative than G37R (*ΔΔG*(*ΔGWT* – *ΔG_mutant_*))suggesting that mutation at 113^th^ position destabilized the protein more in comparison to 37^th^ position (Fig. S13). The vibrational entropy change (*ΔΔS*) in I113T was found to be +0.188 kcal mol^−1^K^−1^ while −0.404 kcal mol^−1^K^−1^ was found for G37R indicating the increase in flexibility for I113T and decrease in flexibility for G37R(Fig.S13). Thus the increase in alpha helical content in I113T mutant arises due to the increase in flexibility in the membrane interacting loop regions as suggested by the DynaMut server. Subsequently, we used ThT fluorescence to probe the aggregation kinetics of G37R and I113T in absence and presence of DPPC SUVs. Results in absence of membrane showed significantly higher aggregation for I113T when compared to G37R. A notable increase in the rate and extent of aggregation was observed for I113T under membrane environment, whereas for G37R, the change was not significant (Fig. 5d). AFM imaging showed linear fibrillar aggregate for I113T in absence of membrane (average length is ~1.5 μm and height is 26 nm) whereas fibrilar network was observed when aggregation occurred in presence of DPPC SUVs (average height is 3.2 nm) (Fig.5e,f). In contrast, small nonfibrilar aggregates were found for G37R mutant both in the absence and presence of membranes (Fig. S14). Finally, we studied the toxic effects of the aggregates of G37R and I113T, which formed in absence and presence of membrane. Using, MTT, we showed that I113T aggregates in absence of membrane were less toxic than I113T aggregates in presence of membrane (Fig. S15). G37R exhibited minimal toxicity both in the absence or presence of membrane (Fig. S15). We observed for these mutants a nice correlation between the extent of calcein release and the populations of nonplanar kinkogtg conformers (Fig. S16 a-f).

**Figure 5:**
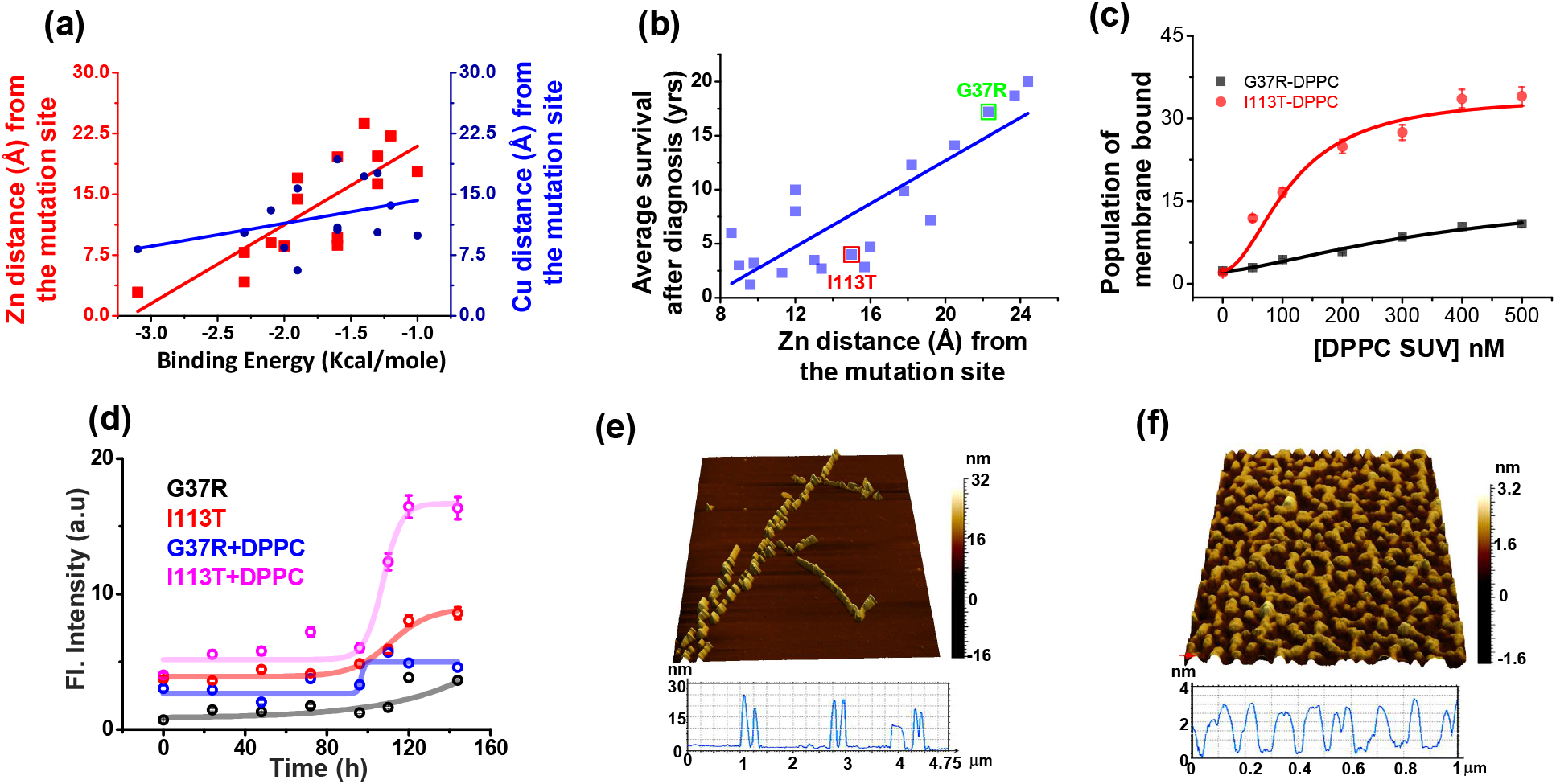
Validation in ALS disease mutants. (a) Computational validation. The distances of the disease mutation sites from the Zn (red) and Cu (blue) co-factor were plotted against the binding energy towards membrane, which were computed. The binding energy decreased linearly as the distance between Zn and mutation site increased (red curve); while there was no significant change for Cu (blue curve). The distance information for these mutants was calculated from their solved structures. (b) The ALS disease severity in terms of average survival time after diagnosis has been plotted against the distance parameters of the mutation points from Zn centre for the disease mutants.(c) percentage populations of membrane bound alexa-labeled protein variants which were obtained from the two component diffusion model fitting of the FCS data for G37R and I113T were plotted against the concentrations of DPPC SUVs added to evaluate the binding affinities of the protein variants towards membranes. (d) Fibril formation kinetics. ThT fluorescence spectra of G37R and I113T were plotted in the absence and presence of DPPC SUVs. The 3D AFM images of I113T in absence (e) and presence (f) of membranes.

## Discussion

### The cofactor derived membrane association model

In this work, we effectively bring a large number of studies together in a coherent framework to address different roles of two metal cofactors (Cu and Zn) and membrane binding on the aggregation and induced toxicity of SOD1. Based on steady state fluorescence, acrylamide quenching and FT-IR data, the investigated proteins could be classified as a) WT and WT-like mutant, H121F and b) apo and apo like mutant, H72F. The battery of experimental approaches clearly validates the prediction from the bWSME model that Zn is largely responsible for the conformational stability of SOD1. The insertion of Zn at the loop IV region stabilizes the loop while reducing the aggregation propensity of the apo protein. We believe that the stabilization of this long loop may play an important role in SOD1 aggregation biology and its relevance in ALS. It may be noted that shortening of the loop has been shown to increase the stability of the protein^45^. This is also important to point out that SOD1 variants from thermophiles (like SOD5 from C. albicans SOD5) have shorter electrostatic loops compared to human SOD1^46^.

In contrast, the absence of Cu does not seem to have any significant effect on the stability of the protein, and the Zn containing Cu deficient protein behaves like WT. We derive a possible maturation/aggregation landscape of SOD1 in the presence of membrane employing data from the current work and available literature (Scheme 1). Since all experiments presented here were carried out using the reduced protein, this scheme does not take dimerization into consideration. SOD1 in its metal free state (apo protein) has been shown to be flexible and inherently dynamic. The sequence sites (48-80 and 120-140) of low folding probability predicted by bWSME calculation (Fig. 1b) has significant overlap with the sequence sites (45-70 and 125-142) of high membrane binding as calculated by OPM(Fig. 2e). It has been found that truncation at Leu126 of C-terminus resulted in a protein with several transmembrane helices^47^. We found that the apo protein has high membrane binding possibility which is directly supported by membrane binding data (Fig. S5b,c). Membrane bound apoSOD1 with an optimized orientation is a requisite for the Cu insertion to take place. Since biological systems have very low free Cu salt^48^, Cu coordination to SOD1 occurs through Cu chaperone protein (CCS) which transfers Cu to apo SOD1 using membrane as a scaffold^49^. Membrane scaffolding decreases the directionality of metal transport compared to a three dimensional search of the metal ions. The domain I of CCS binds Cu through two cysteine residues. Since Cu binding affinity of CCS is less than that of SOD1 (which recruits three or four cysteine residues), Cu transport occurs from CCS to SOD1, and not the other way^50^. Zn binding, which occurs in the next step, stabilizes the loop regions. This event has a few important consequences. First, it has been shown that both CCS and SOD1 exhibit binding to Zn^51^. While the absence of Zn favors heterodimer formation with SOD1 (CCS-SOD1), the presence of Zn facilitates the homodimer (CCS-CCS) configuration. Second, as shown in this paper, Zn coordinated protein has little or minimum membrane binding and hence Zn bound SOD1 is removed from bi-layer. The third consequence comes from the reduced Zn affinity (which may come from a Zn compromised mutation or other reason for the sporadic forms of the disease), which would result in a misfolded membrane bound protein that is aggregation prone and potentially toxic.

As evident, the proposed scheme is Zn centric in which Cu coordination plays minimum role in the disease process. Partial support of this comes directly from the presented data of the distance dependence between ALS mutations sites and Cu/Zn (Figure 5a). Also, the mutant proteins without Cu (but with Zn site coordinated, e.g H121F or G37R) do not show aggregation, neither they induce toxicity as studied by our assay systems. Earlier works have also shown that Cu binding has no effect on SOD1 folding^52^. More importantly, it has been shown that CCS knockout does not affect the onset of the disease or the life span in SOD1 transgenic mice^53^.

It may be important to consider that, although more than 140 SOD1 mutants are known in ALS, the disease is predominantly sporadic. Nevertheless, the disease initiation may still be done by a metal free apo (or Zn free) configuration, which can be generated from the WT protein as a result of a different trigger (and not by a genetic factor). The results obtained from two ALS mutants (I113T and G37R) can be discussed in this context. The present data show that I113T is similar to a zinc deficient protein, with high affinity towards membrane and higher aggregation propensity. In contrast, G37R behaves more like a WT protein. A comparison between the disease phenotype show that I113T is the second most common ALS mutant with average survival of 4.3 years, while for G37R, the average survival increases to 17 years. These results are in excellent correlation with the presented scheme.

Another interesting correlation of the presented scheme comes from a result that Zn supplement with a moderate dose of 60mgkg^−1^day^−1^ can increase the days of survival in a transgenic mouse experiment^54^. Another approach could be using small molecules which would compete with the metal free protein towards membrane binding, an approach used recently in a Parkinson’s disease model^55^. While any drug development initiative targeting ALS and other neurodegenerative diseases suffer from the complications of diseases heterogeneity, poorly understood molecular mechanism and complex delivery avenues, a collaborative and integrative effort involving science and clinic may be needed to find a successful solution.

### The membrane connection

The molecular mechanism associated with ALS induced toxicity has been extensively studied and widely debated^56^. SOD1 aggregates have been found in ALS patient samples^57, 58^. Mutant SOD1 aggregates have been shown to transfer from cell to cell using a prion like propagation mechanism^59^. It has also been shown that the aggregation induced toxicity of SOD1 variants can occur through its attachment on mitochondrial membrane surface in transgenic ALS mice^60^. In parallel, pore formation at the membrane and abnormal ion mobility have been found with SOD1 aggregation^14, 61^. Although the above reports directly link a membrane connection with SOD1 aggregation (and presumably with ALS)-unlike other neurodegenerative diseases like AD and PD, membrane induced aggregation studies with SOD1 have been limited^14^.

Shahmoradian and colleagues reported that the structure of Lewy bodies in Parkinson’s disease consists of α-synuclein and lipid vesicle clusters instead of the long-assumed amyloid fibril core^62^. Using correlative light and electron microscopy (CLEM), they show that the vast majority of Lewy bodies actually consist of clusters of various membranous compartments, instead of amyloid fibrils as previously assumed. Assembly of synaptic vesicles has been shown recently by Fusco et al, in which two different regions of a protein molecule can be used as a ‘double anchor’ to induce the assembly^37^. We think three particular finding can be discussed in this connection. First, the presented membrane deformation assay using GUV clearly show the formation of vesicles assemblies (Fig. 4f) induced by apo_aggm_ and H72F_aggm_. Second, for both protein variants, relatively small aggregates of the proteins (formed in the presence of the membrane) and not the large fibrilar network (formed in the absence of membrane) favored vesicles assemblies. Third, the smaller sized aggregates formed in the presence of membrane seemed to induce more toxicity than the fibrils (formed in the absence). From the above three considerations, it is easy to envision that the anchoring ability of the aggregates towards vesicles assembly would be more efficient for a small-sized aggregates when compared to a fibril. There is an increased interest in the role of the number and nature of membrane lipids in shaping the aggregation landscape of different proteins in neurodegenerative diseases. Given the prion-like propagation in these diseases, our work underscores how characterization of protein-lipid interactions could enhance our molecular understanding of cellular toxicity and enable the identification of therapeutic molecules to mitigate the damage.

## Supporting information

The complete materials and method section has been described in the supporting information file. Additional data (Table S1-S5 and Figure S1-S16) supporting the findings of this study are available in the supplementary information file.

## Acknowledgement

Author A. Sannigrahi acknowledges University Grant Commission, Govt. of India for providing senior research fellowship. S. Karmakar acknowledges the financial support from DBT-funded research project (BT/PR8475/BRB/10/1248/2013). K Chattopadhyay acknowledges funding from the funding from DST SERB (EMR/2016/000310). We thank the director, CSIR-IICB for help and encouragements.

---

**Scheme 1:**
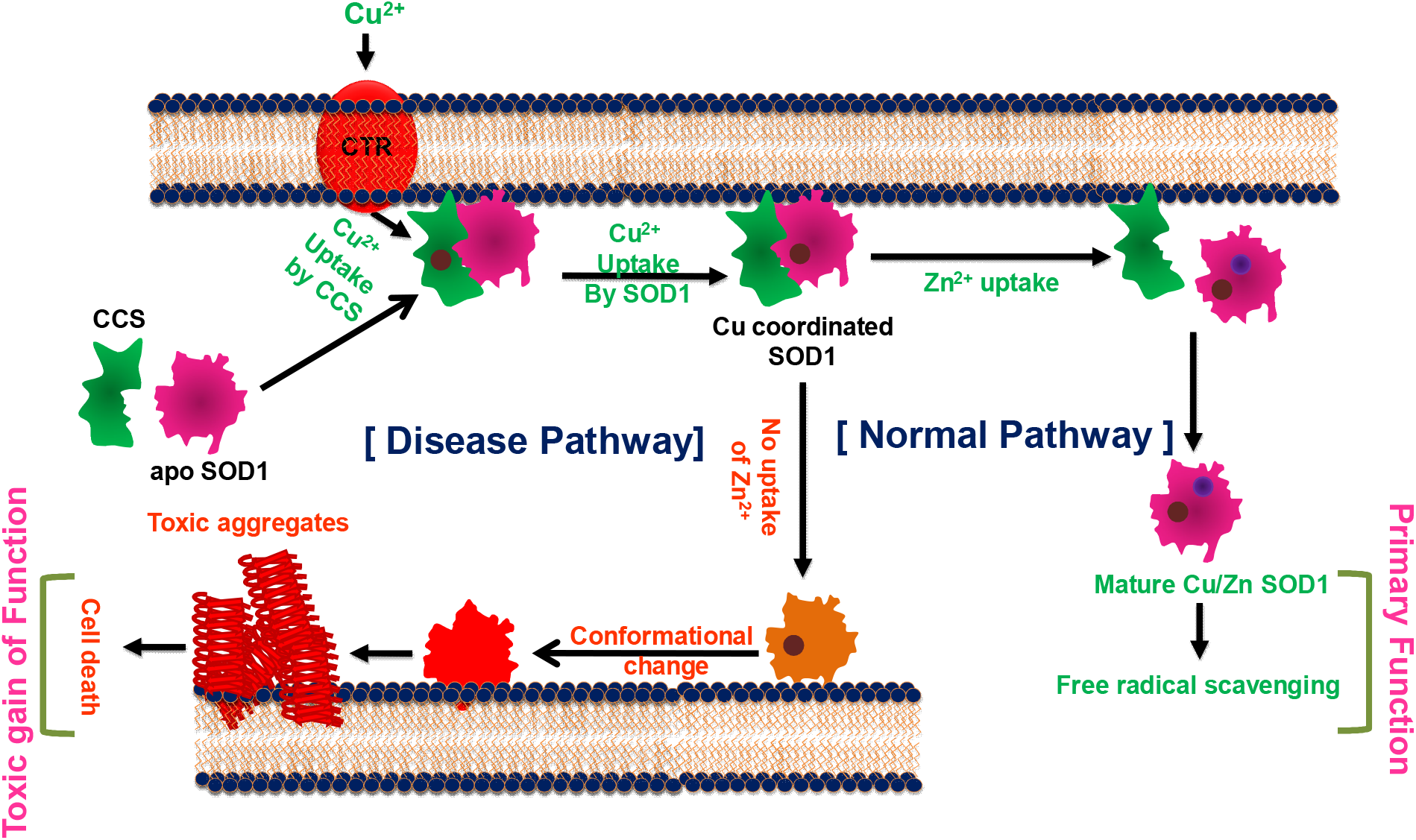
The cofactor derived membrane association model of SOD1 primary and gain of function.

## References

1. Shaw BF, Valentine JS. How do ALS-associated mutations in superoxide dismutase 1 promote aggregation of the protein? Trends in biochemical sciences 32, 78–85 (2007).

2. Bush AI. Is ALS caused by an altered oxidative activity of mutant superoxide dismutase? Nature neuroscience 5, 919–919 (2002).

3. Niwa J-i, et al. Disulfide bond mediates aggregation, toxicity, and ubiquitylation of familial amyotrophic lateral sclerosis-linked mutant SOD1. Journal of Biological Chemistry 282, 28087–28095 (2007).

4. Rodriguez JA, et al. Destabilization of apoprotein is insufficient to explain Cu, Zn-superoxide dismutase-linked ALS pathogenesis. Proceedings of the National Academy of Sciences of the United States of America 102, 10516–10521 (2005).

5. Pasinelli P, et al. Amyotrophic lateral sclerosis-associated SOD1 mutant proteins bind and aggregate with Bcl-2 in spinal cord mitochondria. Neuron 43, 19–30 (2004).

6. Tomik B, et al. Does apoptosis occur in amyotrophic lateral sclerosis? TUNEL experience from human amyotrophic lateral sclerosis (ALS) tissues. Folia neuropathologica 43, 75–80 (2005).

7. Culotta VC, Klomp LW, Strain J, Casareno RLB, Krems B, Gitlin JD. The copper chaperone for superoxide dismutase. Journal of Biological Chemistry 272, 23469–23472 (1997).

8. Mondola P, Damiano S, Sasso A, Santillo M. The cu, Zn superoxide dismutase: not only a dismutase enzyme. Frontiers in physiology 7, 594 (2016).

9. Ilieva H, Polymenidou M, Cleveland DW. Non–cell autonomous toxicity in neurodegenerative disorders: ALS and beyond. The Journal of cell biology 187, 761–772 (2009).

10. Manfredi G, Kawamata H. Mitochondria and endoplasmic reticulum crosstalk in amyotrophic lateral sclerosis. Neurobiology of disease 90, 35–42 (2016).

11. Chng CP, Strange RW. Lipid-associated aggregate formation of superoxide dismutase-1 is initiated by membrane-targeting loops. Proteins: Structure, Function, and Bioinformatics 82, 3194–3209 (2014).

12. Hervias I, Beal MF, Manfredi G. Mitochondrial dysfunction and amyotrophic lateral sclerosis. Muscle & nerve 33, 598–608 (2006).

13. Yamanaka K, et al. Mutant SOD1 in cell types other than motor neurons and oligodendrocytes accelerates onset of disease in ALS mice. Proceedings of the National Academy of Sciences 105, 7594–7599 (2008).

14. Choi I, et al. Lipid molecules induce the cytotoxic aggregation of Cu/Zn superoxide dismutase with structurally disordered regions. Biochimica et Biophysica Acta (BBA)-Molecular Basis of Disease 1812, 41–48 (2011).

15. Wako H, Saitô N. Statistical mechanical theory of the protein conformation. II. Folding pathway for protein. Journal of the Physical Society of Japan 44, 1939–1945 (1978).

16. Muñoz V, Eaton WA. A simple model for calculating the kinetics of protein folding from threedimensional structures. Proceedings of the National Academy of Sciences 96, 11311–11316 (1999).

17. Gopi S, Aranganathan A, Naganathan AN. Thermodynamics and folding landscapes of large proteins from a statistical mechanical model. Current Research in Structural Biology 1, 6–12 (2019).

18. Naganathan AN. Predictions from an Ising-like statistical mechanical model on the dynamic and thermodynamic effects of protein surface electrostatics. Journal of chemical theory and computation 8, 4646–4656 (2012).

19. Rajasekaran N, Gopi S, Narayan A, Naganathan AN. Quantifying protein disorder through measures of excess conformational entropy. The Journal of Physical Chemistry B 120, 4341–4350 (2016).

20. Leinartaite L, Saraboji K, Nordlund A, Logan DT, Oliveberg M. Folding catalysis by transient coordination of Zn2+ to the Cu ligands of the ALS-associated enzyme Cu/Zn superoxide dismutase 1. Journal of the American Chemical Society 132, 13495–13504 (2010).

21. Mojumdar SS, et al. Partially native intermediates mediate misfolding of SOD1 in singlemolecule folding trajectories. Nature communications 8, 1–11 (2017).

22. Chowdhury S, Sen S, Banerjee A, Uversky VN, Maulik U, Chattopadhyay K. Network mapping of the conformational heterogeneity of SOD1 by deploying statistical cluster analysis of FTIR spectra. Cellular and Molecular Life Sciences 76, 4145–4154 (2019).

23. Muneeswaran G, Kartheeswaran S, Muthukumar K, Dharmaraj CD, Karunakaran C. Comparative structural and conformational studies on H43R and W32F mutants of copper–zinc superoxide dismutase by molecular dynamics simulation. Biophysical chemistry 185, 70–78 (2014).

24. Taylor DM, Gibbs BF, Kabashi E, Minotti S, Durham HD, Agar JN. Tryptophan 32 potentiates aggregation and cytotoxicity of a copper/zinc superoxide dismutase mutant associated with familial amyotrophic lateral sclerosis. Journal of Biological Chemistry 282, 16329–16335 (2007).

25. Banci L, et al. Metal-free superoxide dismutase forms soluble oligomers under physiological conditions: a possible general mechanism for familial ALS. Proceedings of the National Academy of Sciences 104, 11263–11267 (2007).

26. Danielsson J, et al. Global structural motions from the strain of a single hydrogen bond. Proceedings of the National Academy of Sciences 110, 3829–3834 (2013).

27. Lomize MA, Pogozheva ID, Joo H, Mosberg HI, Lomize AL. OPM database and PPM web server: resources for positioning of proteins in membranes. Nucleic acids research 40, D370–D376 (2012).

28. Chattopadhyay K, Saffarian S, Elson EL, Frieden C. Measurement of microsecond dynamic motion in the intestinal fatty acid binding protein by using fluorescence correlation spectroscopy. Proceedings of the National Academy of Sciences 99, 14171–14176 (2002).

29. Watanabe S, et al. Mitochondria-associated membrane collapse is a common pathomechanism in SIGMAR1-and SOD1-linked ALS. EMBO molecular medicine 8, 1421–1437 (2016).

30. Casas C, Manzano R, Vaz R, Osta R, Brites D. Synaptic failure: focus in an integrative view of ALS. Brain plasticity 1, 159–175 (2016).

31. Song Y. Synaptic Actions of Amyotrophic Lateral Sclerosis-Associated G85R-SOD1 in the Squid Giant Synapse. Eneuro 7, (2020).

32. Yerbury JJ. Protein aggregates stimulate macropinocytosis facilitating their propagation. Prion 10, 119–126 (2016).

33. McAlary L, Plotkin SS, Yerbury JJ, Cashman N. Prion-Like Propagation of Protein Misfolding and Aggregation in Amyotrophic Lateral Sclerosis. Frontiers in molecular neuroscience 12, 262 (2019).

34. Gohil VM, Greenberg ML. Mitochondrial membrane biogenesis: phospholipids and proteins go hand in hand. Journal of Cell Biology 184, 469–472 (2009).

35. Sannigrahi A, et al. Conformational switch driven membrane pore formation by Mycobacterium secretory protein MPT63 induces macrophage cell death. ACS chemical biology 14, 1601–1610 (2019).

36. Wang J, et al. An ALS-linked mutant SOD1 produces a locomotor defect associated with aggregation and synaptic dysfunction when expressed in neurons of Caenorhabditis elegans. PLoS genetics 5, (2009).

37. Fusco G, et al. Structural basis of synaptic vesicle assembly promoted by α-synuclein. Nature communications 7, 1–12 (2016).

38. Lewis RN, McElhaney RN. Membrane lipid phase transitions and phase organization studied by Fourier transform infrared spectroscopy. Biochimica et Biophysica Acta (BBA)-Biomembranes 1828, 2347–2358 (2013).

39. Maroncelli M, Qi SP, Strauss HL, Snyder RG. Nonplanar conformers and the phase behavior of solid n-alkanes. Journal of the American Chemical Society 104, 6237–6247 (1982).

40. Wang Q, Johnson JL, Agar NY, Agar JN. Protein aggregation and protein instability govern familial amyotrophic lateral sclerosis patient survival. PLoS biology 6, (2008).

41. Milardi D, Pappalardo M, Grasso DM, La Rosa C. Unveiling the unfolding pathway of FALS associated G37R SOD1 mutant: a computational study. Molecular BioSystems 6, 1032–1039 (2010).

42. Krishnan U, Son M, Rajendran B, Elliott JL. Novel mutations that enhance or repress the aggregation potential of SOD1. Molecular and cellular biochemistry 287, 201–211 (2006).

43. Banci L, et al. Structural and dynamic aspects related to oligomerization of apo SOD1 and its mutants. Proceedings of the National Academy of Sciences 106, 6980–6985 (2009).

44. Rodrigues CH, Pires DE, Ascher DB. DynaMut: predicting the impact of mutations on protein conformation, flexibility and stability. Nucleic acids research 46, W350–W355 (2018).

45. Yang F, Wang H, Logan DT, Mu X, Danielsson J, Oliveberg M. The cost of long catalytic loops in folding and stability of the ALS-associated protein SOD1. Journal of the American Chemical Society 140, 16570–16579 (2018).

46. Gleason JE, et al. Candida albicans SOD5 represents the prototype of an unprecedented class of Cu-only superoxide dismutases required for pathogen defense. Proceedings of the National Academy of Sciences 111, 5866–5871 (2014).

47. Lim L, Lee X, Song J. Mechanism for transforming cytosolic SOD1 into integral membrane proteins of organelles by ALS-causing mutations. Biochimica et Biophysica Acta (BBA)-Biomembranes 1848, 1–7 (2015).

48. Rae T, Schmidt P, Pufahl R, Culotta V, O’Halloran TV. Undetectable intracellular free copper: the requirement of a copper chaperone for superoxide dismutase. Science 284, 805–808 (1999).

49. Pope CR, De Feo CJ, Unger VM. Cellular distribution of copper to superoxide dismutase involves scaffolding by membranes. Proceedings of the National Academy of Sciences 110, 20491–20496 (2013).

50. Banci L, Bertini I, Ciofi-Baffoni S, Kozyreva T, Zovo K, Palumaa P. Affinity gradients drive copper to cellular destinations. Nature 465, 645–648 (2010).

51. Proescher JBG. Effects of the CCS copper chaperone on SOD1 mutants linked to amyotrophic lateral sclerosis. The Johns Hopkins University (2008).

52. Bruns CK, Kopito RR. Impaired post-translational folding of familial ALS-linked Cu, Zn superoxide dismutase mutants. The EMBO journal 26, 855–866 (2007).

53. Subramaniam JR, et al. Mutant SOD1 causes motor neuron disease independent of copper chaperone–mediated copper loading. Nature neuroscience 5, 301–307 (2002).

54. Ermilova IP, Ermilov VB, Levy M, Ho E, Pereira C, Beckman JS. Protection by dietary zinc in ALS mutant G93A SOD transgenic mice. Neuroscience letters 379, 42–46 (2005).

55. Fonseca-Ornelas L, et al. Small molecule-mediated stabilization of vesicle-associated helical α-synuclein inhibits pathogenic misfolding and aggregation. Nature communications 5, 1–11 (2014).

56. Broom HR, et al. Folding and aggregation of Cu, Zn-superoxide dismutase. In: Amyotrophic Lateral Sclerosis). InTech (2012).

57. Gruzman A, et al. Common molecular signature in SOD1 for both sporadic and familial amyotrophic lateral sclerosis. Proceedings of the National Academy of Sciences 104, 12524–12529 (2007).

58. Guareschi S, et al. An over-oxidized form of superoxide dismutase found in sporadic amyotrophic lateral sclerosis with bulbar onset shares a toxic mechanism with mutant SOD1. Proceedings of the National Academy of Sciences 109, 5074–5079 (2012).

59. Münch C, O’Brien J, Bertolotti A. Prion-like propagation of mutant superoxide dismutase-1 misfolding in neuronal cells. Proceedings of the National Academy of Sciences 108, 3548–3553 (2011).

60. Zhai J, et al. Proteomic characterization of lipid raft proteins in amyotrophic lateral sclerosis mouse spinal cord. The FEBS journal 276, 3308–3323 (2009).

61. Ray SS, Nowak RJ, Strokovich K, Brown Jr RH, Walz T, Lansbury Jr PT. An intersubunit disulfide bond prevents in vitro aggregation of a superoxide dismutase-1 mutant linked to familial amytrophic lateral sclerosis. Biochemistry 43, 4899–4905 (2004).

62. Shahmoradian SH, et al. Lewy pathology in Parkinson’s disease consists of crowded organelles and lipid membranes. Nature neuroscience 22, 1099–1109 (2019).

