## Supporting Information for "The metal cofactor zinc and interacting membranes modulate SOD1 conformation-aggregation landscape in an *in vitro* ALS Model"

**Materials.** 1,2-dipalmitoyl-sn-glycero-3-phosphocholine (DPPC), 1,2-dioleoyl-sn-glycero-3-phosphoethanolamine (DOPE), phosphoinositol (PI),1,2-dioleoyl-sn-glycero-3-phospho-L-serine (DOPS) and cardiolipin(CL) were purchased from Avanti Polar Lipids Inc. (Alabaster, AL, USA). 1,1'-dioctadecyl-3,3,3',3'-tetramethylindotricarbocyanine iodide (DiIC-18(3)) was purchased from Invitrogen (Eugene, Oregon, USA). All other necessary chemicals were obtained from Aldrich (St. Louis, USA) and Merck (Mumbai, India).

**Experimental Methods**

**Expression and Purification of SOD1.** Recombinant SOD1 was over-expressed in *E. coli* (BL21 DE3 strain). The over-expression of SOD1 was induced with 1 mM IPTG. The induction was coupled with metalation where 1mM $CuSO_4$ was added directly to the Luria-Bertani culture media so as to ensure proper metal loading over the protein. Followed by induction, the cells were allowed to grow for 3.5 hours. The cells were pelleted down by centrifuging at 6000 rpm for 15 minutes at 4 °C followed by re-suspension in pre-chilled lysis buffer (20 mM Tris-HCl + 500 mM NaCl, pH 8.0). After thorough re-suspension in lysis buffer the cells were subjected to sonication (20 pulses, each of 30 seconds pulse time and an interim time frame of 1 minute). Unbroken cells and debris were removed by another act of centrifugation at 10,000 rpm for 10 minutes. The soluble fraction obtained thereafter was carefully removed and allowed to bind to Ni-NTA agarose resin. The Ni-NTA column was washed with 40 ml wash buffer (20 mM Tris-HCl, 500 mM NaCl and 50 mM imidazole, pH 8.0) followed by elution with 20 mM Tris HCl, 500 mM NaCl and 500 mM imidazole, pH 8.0. The eluted fractions were pooled according to their tentative protein content as per their absorbance at 280 nm. The post elution fractions were subjected to dialysis in 20 mM Na-phosphate buffer pH 7.5. In all our protein concentration measurements ultraviolet spectroscopy was deployed and SOD1 concentration was determined by considering the monomeric molar extinction coefficient of 5,500 $M^{-1}$ $cm^{-1}$ at 280 nm.[1] The identity of the protein was confirmed by SDS PAGE and MS/MS. The metal content was confirmed by the Atomic Absorption Spectroscopy and activity assays as reported earlier.[2]

**Site-Directed Mutagenesis.** The recombinant plasmid pET-19b, containing the gene for hSOD1 with a poly Histidine tag at the N-terminal end, was used as a template for mutagenesis using the QuickChange XL Site-Directed Mutagenesis Kit (Stratagene, Agilent, USA). The mutagenic primers containing the mutations (shown in bold type) for the replacement of His-72 with a

Phenylalanine residue (H72F) and His-121 with a Phenylalanine residue (H121F) are indicated below. Both primers were annealed to the same target sequence on opposite strands of pET-19b. The site-directed mutagenesis was performed using protocol as described by the manufacturer. The clone used for production of the H72F and H121F mutant was confirmed by DNA sequencing. Proteins were expressed in BL21 (DE3) pLysS*E. coli* cells by induction with 0.5 mM isopropyl-1- thio-*β*-D-galactopyranoside (IPTG) at 37 °C for 4 h. Cells were resuspended in ice cold 20 mM Tris-HCl, 500 mM NaCl, pH 8.0, containing protease inhibitor (2 mM PMSF) and lysed by sonication. Unbroken cells and debris were removed by centrifugation at 10000*g* for 10 min. Binding of soluble proteins from the supernatant to Ni-NTA agarose resin (Qiagen) was done overnight. The Ni-NTA flow through was collected for analysis. The Ni-NTA resin was washed with 40 ml of wash buffer (20 mM Tris-HCl pH 8, 500 mM NaCl, 50 mM imidazole). Elution was done with wash buffer containing 500 mM imidazole. The designed primers for different mutants are as follows:

| | |
|---|---|
| H72F | 5'-cctcactttaatcctctatccagaaaattcggtgggccaaagg-3' |
| H72F_antisense | 5'-cctttggcccaccgaattttctggatagaggattaaagtgagg-3' |
| H121F | 5'-ggccgcacactggtggtctttgaaaaagcagatgactt-3' |
| H121F_antisense | 5'-aagtcatctgctttttcaaagaccaccagtgtgcggcc-3' |

**Preparation of apo SOD1**. Preparation of apo enzyme from WT SOD1 by metal chelation was carried out following the earlier reported protocol.[3] Overnight dialysis of WT SOD1 in 50 mM Na-acetate, 10 mM EDTA, pH 3.8 was done so as to ensure proper removal of metal ions. EDTA was removed by repeated dialysis in 50 mM Na-acetate, pH 5.2 and in 20 mM Na-phosphate, pH 7.5. The de-metalation was ensured by an activity assay of SOD1 indexing photo-oxidation of pyrogallol at pH 8.

**Enzyme assay for SOD1.** Inhibition of superoxide anion ($O_2^-$) mediated pyrogallol auto-oxidation in alkaline buffer (Tris-cacodylate, pH 8.2) by SOD1 was performed for measuring the activity of the metalloenzyme.[2] Absorption of the oxidized product at 420 nm was monitored with respect to time to assess the activity of Cu/Zn SOD1. Extent of pyrogallol auto-oxidation was measured from the ratio of product absorption at 420 nm in absence and presence of SOD1. Time plot of 0.2 mM pyrogallol was constructed. Then protein (100 nM) was added and time plot was recorded. Product formation in absence of SOD1 was taken as the reference value of 100.

**Preparation of Small Unilamellar Vesicles (SUVs).** The appropriate amount of lipids in chloroform (concentration of stock solution is 25 mg $mL^{-1}$) was transferred to a 10 mL glass bottle. The organic solvent was removed by gently passing dry nitrogen gas. The sample was then placed in a desiccator connected to a vacuum pump for a couple of hours to remove traces of the leftover solvent. A required volume of 20 mM sodium phosphate buffer at pH 7.4 was added to the dried lipid film so that the final desired concentration (10 mM) was obtained. The lipid film with the buffer was kept overnight at 4°C to ensure efficient hydration of the phospholipid heads. Vortexing of the hydrated lipid film for about 30 min. produced multilamellar vesicles (MLVs). Long time vortexing was occasionally required to make uniform lipid mixtures. This MLV was used for optical clearance assay and Dynamic Light Scattering (DLS) experiment. For preparing the SUV, as-formed MLV was sonicated using a probe sonicator at an amplitude 45% for 30 mins. and after that sample was centrifuged at 5000 rpm to sediment the tungsten artifacts and finally it was filtered by 0.22 µm filter unit. Size of the vesicles was measured by DLS to be ~ 70 nm in diameter.

**Tryptophan Quenching Experiments.** Steady state fluorescence spectroscopy and acrylamide quenching measurements in free and in membrane bound conditions were carried out using a PTI fluorimeter (Photon Technology International, USA). A cuvette with 1cm path length was used for the fluorescence measurements. For the tryptophan fluorescence quenching experiments, an excitation wavelength of 295 nm was used to eliminate the contributions from tyrosine fluorescence. Fluorescence data were recorded using a step size of 1 nm and an integration time of 1sec.Excitation and emission slits were kept at 5 nm in each case. Emission spectra between 305nm and 450 nm were recorded in triplicate for each experiment. Typical protein concentration of 10 µM was used for each quenching experiment and 1:200 protein-lipid molar ratio was maintained. The protein solutions were incubated at room temperature for 1 hour and then titrated using a stock of 10M acrylamide. Necessary background corrections and inner filter effect corrections were made for each experiment.

**Acrylamide Quenching Data Analysis**. Assuming $I$ and $I_o$ are the tryptophan fluorescence intensity of the proteins in the presence and absence of acrylamide concentration [$Q$], the Stern-Volmer Equation[4] can be represented as follows:

$$\frac{I_0}{I} = 1 + K_{sv}[Q] \quad (1)$$

Where, $K_{sv}$ is the Stern-Volmer constant, which can be determined from the slope of the linear plot of $I_0/I$ *versus* acrylamide concentrations [$Q$].

**Fourier Transform Infrared Spectroscopy (FTIR).** FTIR spectra of WT SOD1, apo SOD1and mutants in absence and presence of denaturants were acquired using a Bruker 600 series FTIR spectrometer. The FTIR spectral readouts were collected at pH 7.5 immediately after dispensing the proteins in respective buffer solutions. Buffer baseline was subtracted before taking each spectrum. The deconvolution of raw spectra in the amide I region (1700 $cm^{-1}$ to 1600 $cm^{-1}$) was done using least-squares iterative curve fitting to Gaussian/Lorentzian line shapes. The assignment of peaks was done using previously described spectral components associated with different secondary structures.[5] For investigating the morphological changes of bilayer due to interaction of protein variants and preformed aggregates, FTIR spectroscopy was utilized by using typical lipid concentration 2 mM. Background corrections were done for each and every experiment. We specifically evaluated the vibrational changes in the planar DPPC bilayer through the measurement of the changes in $CH_2$ wagging band frequency (1280-1460 $cm^{-1}$) of the hydrocarbon tail region was considered for evaluation.

**AFM Studies.** Aliquots of aggregating samples were withdrawn after prolonged incubation at 37 °C and were diluted with 5 mM phosphate buffer, pH 7.4. A 5–8 µl aliquot was taken from the diluted sample and deposited on freshly cleaved mica for 10 min. The typical protein-concentration was taken 500 nM. After removing the excess liquid, the aggregates were rinsed with MilliQ water and then dried with a stream of nitrogen. Images were acquired at ambient temperature using a Bioscope Catalyst AFM (Bruker Corporation, Billerica, MA) with silicon probes. The standard tapping mode was used to image the morphology of aggregates. The nominal spring constant of the cantilever was kept at 20–80 N/m. The spring constant was calibrated by a thermal tuning method. A standard scan rate of 0.5 Hz with 512 samples per line was used for imaging the samples. A single third order flattening of height images with a low pass filter was done followed by section analysis to determine the dimensions of aggregates.

**ThT Fluorescence Assay.** WT SOD1, apo, H72F and H121F proteins were subjected to mechanical agitation at 200 rpm at 37 °C for 350 hours. The protein concentrations for the aggregate preparation were kept 50 µM in 20 mM sodium phosphate buffer at pH 7.5.The protein species were treated with 1.2µM TCEP. For the measurement of membrane induced protein aggregation, WT and other protein variants were incubated in lipid/protein ratio 2:1 under the treatment of TCEP. Aliquots were thereafter subjected to ThT addition and fluorescence measurements were taken using an integration time of 0.3 s. The steady state fluorescence was monitored using an excitation wavelength of 450 nm, and the values of emission intensity at 485 nm were recorded.

**Assay for Permeabilization of Lipid Vesicles.** The ability of protein aggregates to promote the release of calcein from entrapped SUVs composed of DOPC:DOPE:DOPS in the ratio 2:5:3 was checked by monitoring the increase in fluorescence intensity of calcein. Calcein-loaded liposomes were separated from non-encapsulated (free) calcein by gel filtration on a Sephadex G-75 column (Sigma) using an elution buffer of 10 mM MOPS, 150 mM NaCl and 5 mM EDTA (pH 7.4), and lipid concentrations were estimated by complexation with ammonium ferro-thiocyanate[6]. Fluorescence was measured at room temperature (25 $^{\circ}$C) in a PTI spectro-fluorometer using a 1cm path length cuvette. The excitation wavelength was 490 nm and emission was set at 520 nm. Excitation and emission slits with a nominal bandpass of 3 and 5 nm were used, respectively. The high concentration (10 mM) of the entrapped calcein led to self-quenching of its fluorescence resulting in low fluorescence intensity of the vesicles ($I_B$). Release of calcein caused by addition of proteins aggregates led to the dilution of the dye into the medium, which could therefore be monitored by an enhancement of fluorescence intensity ($I_F$). This enhancement of fluorescence is a measure of the extent of vesicle permeabilization. The experiments were normalized relative to the total fluorescence intensity ($I_T$) corresponding to the total release of calcein after complete disruption of all the vesicles by addition of Triton X-100 (2% v/v). The percentage of calcein release in the presence of different aggregates of SOD1 protein species was calculated using the equation[7]:

$$\% \, release = 100 \, \frac{(I_F - I_B)}{(I_T - I_B)} \qquad (2)$$

where, $I_B$ is the background (self-quenched) intensity of calcein encapsulated in vesicles, $I_F$ represents the enhanced fluorescence intensity resulting from the dilution of dye in the medium caused by protein-aggregates induced release of entrapped calcein. $I_T$ is the total fluorescence intensity after complete permeabilization is achieved upon addition of Triton X-100. Typical SUVs concentration was taken 100 μM and 5μM concentration of each protein aggregate sample was used. The pore formation rate constants (^, cm-1) were calculated through the exponential fitting of the growth kinetics due to leakage of calcein from the entrapped SUVs.

**Preparation of GU**Vs. GUVs were formed in 0.1 M Sucrose prepared in 1 mM HEPES (pH 7.4) buffer using electroformation, as described by Pott et al.[8] Briefly, 20 μL of a 1 mM lipid (DOPC:DOPE:PI:DOPS:CL in the ratio 4.5:2.5:1:0.5:1.5) solution in chloroform were spread onto the surfaces of two conductive glasses (coated with Fluor Tin Oxide), which were then placed with their conductive sides facing each other. These droplets were allowed to dry overnight in a closed chamber containing saturated solution of NaCl. This is to avoid complete drying of the droplets. The hydration of these droplets facilitates electroformation process. Electroformation chamber was made using Teflon spacer of thickness ∼2 mm. This electro swelling chamber was filled with 0.1 M sucrose solution and branched to an alternating power generator at 1.5 V and 15 Hz frequency during 2h at room temperature (22–25 °C). The vesicles solution was then carefully transferred to an eppendorf vial and kept at rest at 4°C before use. The average diameter of the GUV obtained was 10−100 μm. For single GUV imaging we selected the GUV of size ~30 μm for eliminating the effect of curvature. GUVs were diluted in 0.1 M glucose, prepared in 1 mM HEPES (pH 7.4), for observation. A typical observation experiment, using an inverted microscope, was made in an observation chamber by mixing 30 μL of the GUV solution with 100 μL of a 0.1 M glucose solution. The slight density difference between the inner and outer solutions drives the vesicles to settle at the bottom of the slide and provides better contrast while observing under phase contrast.

**Phase Contrast Optical Microscopy**. For the microscopy experiment, 5 μM concentrations of protein aggregates of different variants were dissolved in the glucose solution prepared in 1 mM HEPES buffer (pH 7.4) were added to the vesicle solution. Phase contrast microscopy was performed using an inverted microscope (DMi8) from Leica (Wetzlar, Germany). Observation chamber consisted of a glass slide with rubber spacers. The chamber was then closed immediately for observation under phase contrast microscope after placing the samples. Response of individual GUV when exposed to the protein aggregates solution was continuously recorded with time using a CCD camera. Images were analyzed using the image analysis software, Image J. A straight line was drawn across the GUV to obtain an intensity profile. Peak to peak intensity ($I_{ptp}$) across the halo region was calculated. Average $I_{ptp}$ was obtained from several line profiles across the GUV. The time in second versus $I_{ptp}$ was plotted in order to observe any significant change in the intensity profile of the GUV. We have calculated the $I_{ptp}$ values for at least three different GUVs treated with different aggregate species and averaged the values to calculate the deformation rate constant ($\lambda$, $sec^{-1}$).

**Cell Culture and cytotoxicity assay.** Neuroblastoma cell lines SHSY5Y were acquired from the national cell repository (National Centre for Cell Science, Pune, India). Cells were maintained in Dulbecco's Modified Eagle's Media (DMEM) which in turn were supplemented with 10% heat-inactivated Fetal Bovine Serum (FBS) respectively, 4.5 g/L of glucose, 1.5 g/L sodium bicarbonate, 110 mg/L sodium pyruvate, 4 mM L-glutamine, 50 units/ml penicillin G, and 50 μg/ml streptomycin in humidified air containing 5% $CO_2$ at 37°C. Sub-culturing was done by allowing the passaging of cells as per ATCC recommendations (ATCC, Manassus, VA, USA). Cells were cultured in both serum and antibiotic free culture medium before each experiment. MTT assay was directed to evaluate the cell cytotoxicity.[9] For the initial screening experiment, the SHSY5Y cells ($4\times10^3$ cells per well) were seeded in a 96 well plate and left in an incubator followed by treatment with different protein aggregates variants (5μM) for 12 h. After 12 h of incubation, cells were washed with PBS, and then the MTT solution was added to each well and kept in an incubator for 4 h to form formazan salt. Then the formazan salt was solubilized using DMSO and the absorbance was observed at 595 nm using an ELISA reader (Emax, Molecular device, USA).

**Labeling of the Protein with Alexa488Maleimide.** All the SOD1 protein species were labeled with Alexa488Maleimide (Alexa488) using a previously published procedure.[10] Briefly, Alexa488 dissolved in DMSO was slowly added to 2 mg $mL^{-1}$ solution of the protein under constant stirring. The molar ratio between the dye and protein was maintained at 10:1. The reaction mixture was incubated at 4 °C for 5 h, with shaking after every 30 min. The labeling reaction was then stopped by adding excess β-mercaptoethanol. Excess free dye from the reaction mixture was removed by extensive dialysis followed by column chromatography using a Sephadex G20 column which was pre-equilibrated with 20 mM sodium phosphate buffer (pH 7.5).

**Fluorescence Correlation Spectroscopy (FCS) Experiments and Data Analysis.** FCS experiments were carried out using a dual channel ISS Alba V system equipped with a 60X water-immersion objective (NA 1.2). Samples were excited with an argon laser at 488 nm. All protein data were normalized using the $\tau_D$ value obtained with the free dye (Alexa488) which was measured under identical conditions. For a single-component system, diffusion time ($\tau_D$) of a fluorophore and the average number of particles ($N$) in the observation volume can be calculated by fitting the correlation function [$G(\tau)$] to Eq.3 :

$$G(\tau) = 1 + \left(\frac{1}{N\left(1+\frac{\tau}{\tau_D}\right)}\right)\frac{1}{\sqrt{1+S^2\frac{\tau}{\tau_D}}} \quad \text{-------------(3)}$$

where, $S$ is the structure parameter, which is the depth-to-diameter ratio. The characteristic diffusion coefficient ($D$) of the molecule can be calculated from $\tau_D$ using Eq. 4:

$$\tau_D = \frac{\omega^2}{4D} \quad \text{--------------------------------------(4)}$$

where, $\omega$ is the radius of the observation volume, which can be obtained by measuring the $\tau_D$ of a fluorophore with known $D$ value. The value of hydrodynamic radius ($r_H$) of a labelled molecule can be calculated from $D$ using the Stokes-Einstein equation [Eq. 5]:

$$D = \frac{kT}{6\pi\eta r_H} \quad \text{----------------------------------------(5)}$$

where, $k$ is the Boltzmann constant, $T$ is the temperature and $\eta$ corresponds to the viscosity of the solution.[11]

**Wako-Saitô-Muñoz-Eaton (WSME) Model.** We employ the Ising-like WSME model[12, 13] with the block approximation[14] to predict the conformational landscape of SOD1 oxidized monomer and its variants using the PDB structure 4FF9 as the reference. Briefly, the model assigns binary variables of 1 or 0 for folded or unfolded status of residues, respectively. We employ a version of the model that accounts of single-stretches of folded blocks (single-sequence approximation, SSA), two stretches of folded blocks (double-sequence approximation, DSA) and DSA allowing for interactions across the folded islands if they are interacting in the folded structure. The 151-residue protein SOD1 is therefore reduced to a collection of 49 sequential blocks on assuming a block length of 3thus reducing the number of microstates from >42,700,000 (in the residue-level version of the model) to 461,826. The energetics of the model involves van der Waals (vdW) interactions identified with a 6 Å heavy-atom cut-off (with a vdW interaction energy of $\xi$), all-to-all Debye-Hückel electrostatics, and simplified solvation (defined by the heat capacity change per native contact of) [15].Residues identified to be fully folded are assigned an entropic penalty of -13.6 J mol$^{-1}$ K$^{-1}$ per residue ($\Delta S_{conf}$).The apo form of SOD1 is simulated by assigning an excess conformational entropy of -19.7 J mol$^{-1}$ K$^{-1}$per residue[16] for the stretches of residues 49-82 (loop IV, Zn binding loop) and 121-142 (loop VII, Cu binding loop), as reported from NMR order parameter measurements[17]. To simulate order in either one or both the loops the conformational entropy of residues in the loop is modified to -13.6 J mol$^{-1}$ K$^{-1}$ per residue (*i.e.* a lower penalty for folding) thus mimicking the variants of SOD1 (Zn bound, Cu bound, and Holo forms). The van der Waals interaction energies are fixed to -35.9, -38.2, -42.2, and -48.9 J mol$^{-1}$ for the Holo, Zn bound, Cu bound, and apo variants, respectively, to simulate iso-stability conditions at 298 K. The heat capacity change per native contact is fixed to -0.36 J mol$^{-1}$ K$^{-1}$ per native contact. All prolines are assigned an entropic penalty of zero to account for their rigidity. Residue probabilities and folding mechanism as a function of the number of structured blocks are predicted at iso-stability conditions (i.e. a stability 25 kJ mol$^{-1}$at 298 K) following established protocols by accumulating partial partition functions[15, 16].

**OPM (Orientations of Proteins in Membranes).** To gain an insight as to how the WT and metal starved variants of SOD1 interact with membrane, we resorted to computational approaches. Protein orientations in membranes were theoretically calculated by minimizing a protein's transfer energy from water to a planar slab that serves as a crude approximation of the membrane hydrocarbon core. For WT SOD1 we referred to the solved structure 4BCY and for the metal starved forms in vacuo *ab-initio* models were prepared from Zhang Lab server[18]. The membrane binding propensity was calculated submitting the co-ordinate information of the protein forms to OPM server.[19] A protein was considered as a rigid body that freely floats in the planar hydrocarbon core of a lipid bilayer. Accessible surface area is calculated using the subroutine SOLVA from NACCESS with radii of Chothia (1975) and without hydrogen.[20] In OPM, solvation parameters are derived specifically for lipid bilayers and normalized by the effective concentration of water, which changes gradually along the bilayer normal in a relatively narrow region between the lipid head group regions and the hydrocarbon core.

**Table S1** Metal contents (Cu and Zn) in WT and other mutants (H121F, H72F and apo) as obtained from Atomic Absorption Spectroscopy.

| Protein Forms | Cu content | Zn Content |
|---|---|---|
| WT | 4.5μM | 3.9 μM |
| H121F | <1 μM | 3.8 μM |
| H72F | 4.1 μM | <1.2 μM |
| apo | < 0.5 μM | < 0.25 μM |

**Table S2** The values of Stern volmer quenching constants for Trp 32 residue of all the protein variants in absence($K_{sv}$,$M^{-1}$) and presence of DPPC SUVs ($K_{svm,}$ $M^{-1}$) as obtained from acryamide quenching for WT SOD1 and all the mutants including apo SOD1.

| Proteins | $K_{sv}$ | $K_{svm}$ | $K_{sv}$/ $K_{svm}$ |
|---|---|---|---|
| WT SOD1 | 6.8±0.1 | 5.7±0.2 | 1.19 |
| H121F | 8.0±0.1 | 6.3±0.1 | 1.26 |
| H72F | 12.7±0.3 | 7.0±0.2 | 1.82 |
| apo SOD1 | 14.3±0.1 | 7.6±0.2 | 1.88 |

**Table S3** Binding constants ($K_a$,$M^{-1}$) of the protein variants with model DPPC SUVs as obtained from the FCS study for WT SOD1 and all the mutants.

| **Systems** | **Asssociation constants($K_a$,$M^{-1}$)** |
|---|---|
| **WT SOD1+DPPC SUV** | $(4.6 \pm 0.1) x10^6$ |
| **H121F+DPPC SUV** | $(6.6\pm0.2)x10^6$ |
| **H72F+DPPC SUV** | $(9.6\pm0.4)$ $x10^7$ |
| **apo+DPPC SUV** | $(9.8\pm0.1)$ $x10^7$ |
| **G37R-DPPC SUV** | $(2.6\pm0.3)x10^6$ |
| **I113T-DPPC SUV** | $(8.6\pm0.2)x10^6$ |

**Table S4** Log phase mid points of different protein-variants obtained from ThT assay.

| Systems | Log phase mid-point (h) |
|---|---|
| WT SOD1 | Not detectable |
| WT SOD1+DPPC SUV | Not detectable |
| H121F | Not detectable |
| H121F+DPPC SUV | Not detectable |
| H72F | 167.2 |
| H72F+DPPC SUV | 90.8 |
| apo SOD1 | 112.8 |
| apo+DPPC SUV | 55.9 |

**Table S5** SOD1 disease mutants, their corresponding distance parameters in terms of the distances of mutational stress points from the Zn and Cu centre and the membrane binding energies of the disease mutants

| Mutants | Mutational points distances from Zn centre(Å) | Mutational points distances from Cu centre(Å) | Membrane binding energies (ΔG, kcal/mole) |
|---|---|---|---|
| A4V | 19.6 | 19.3 | -1.6 |
| C6A | 17 | 15.7 | -2.1 |
| G37R | 23.7 | 17.2 | -1.0 |
| L38V | 22.2 | 13.6 | -1.2 |
| H43R | 17.8 | 9.9 | -1.0 |
| H46R | 7.8 | 10.2 | -2.3 |
| H80R | 4.2 | 10.3 | -2.3 |
| G85R | 8.6 | 8.4 | -2.0 |
| G93A | 20.5 | 22.7 | -1.8 |
| C111A | 11.9 | 18.3 | -3.3 |
| C112S | 19.7 | 17.6 | -1.3 |
| I113T | 16 | 18.5 | -3.2 |
| D124V | 9 | 13 | -2.1 |
| H72F | 2.9 | 8.2 | -3.1 |
| H121F | 14.4 | 5.6 | -1.9 |
| S134N | 9.6 | 10.9 | -1.6 |
| C147S | 16.3 | 10.3 | -1.7 |

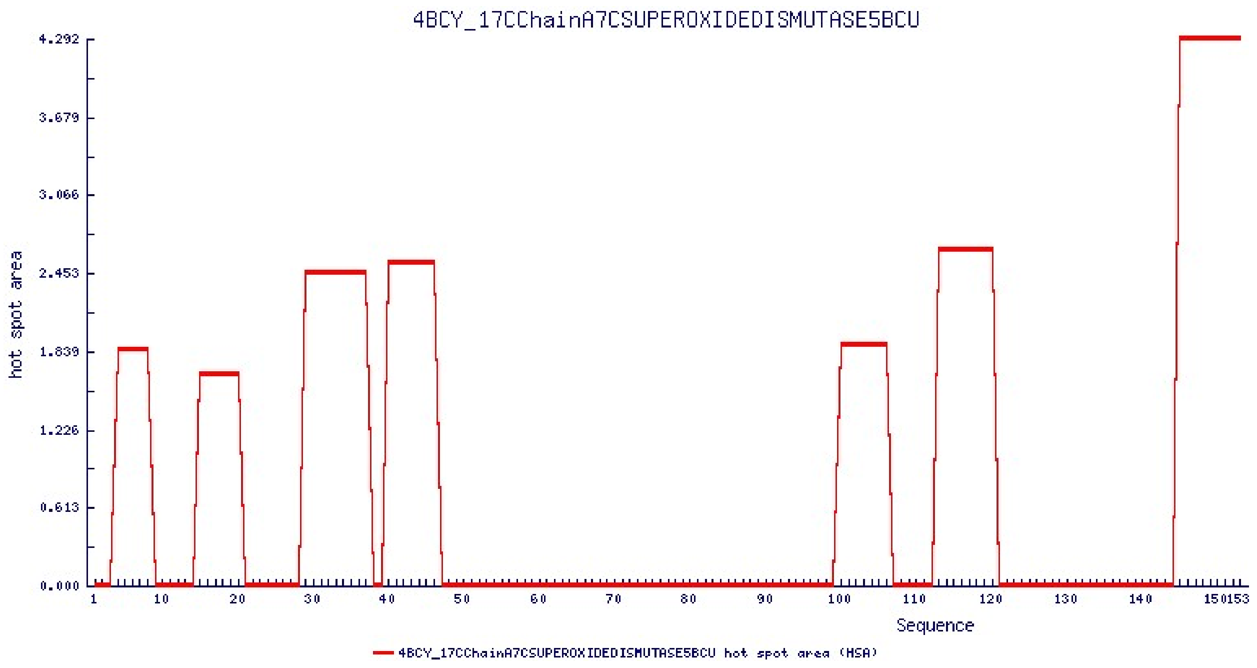


**Figure S1** Aggregation prone regions as a function of sequence from AGGRESCAN software (http://bioinf.uab.es/aggrescan/ ). Results point to the N-terminal residues 1-50 as the hotspot region for aggregation.

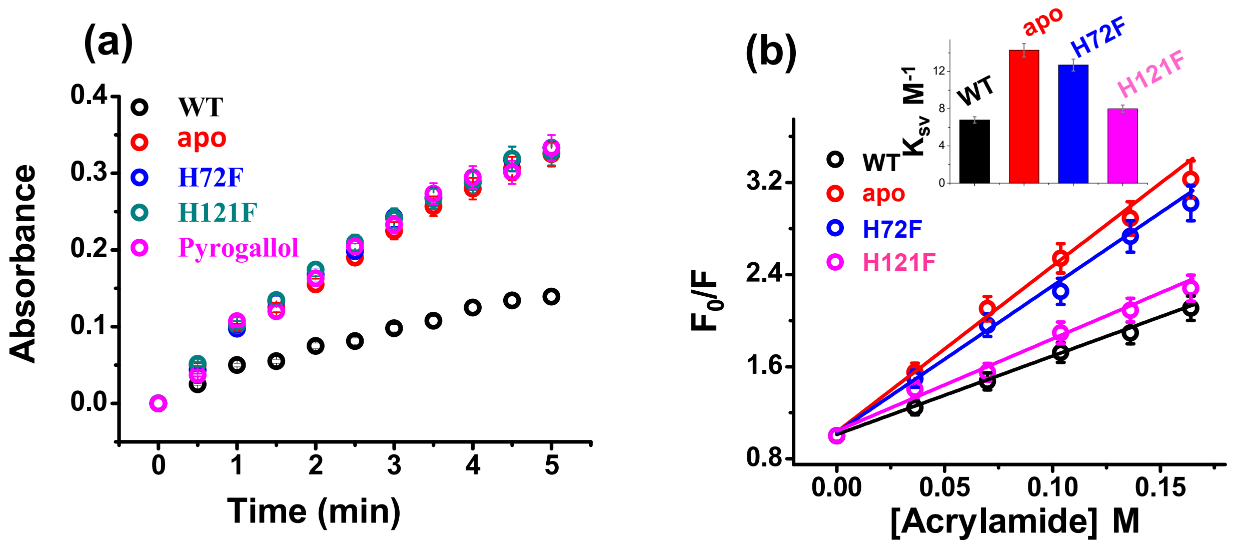


**Figure S2** (a) Activity assay (pyrogallol auto-oxidation) for WT and other metal mutants of SOD1.(b) Acryamide quenching experiments for the different variants. Inset shows the values of $K_{sv}$ of different protein species.

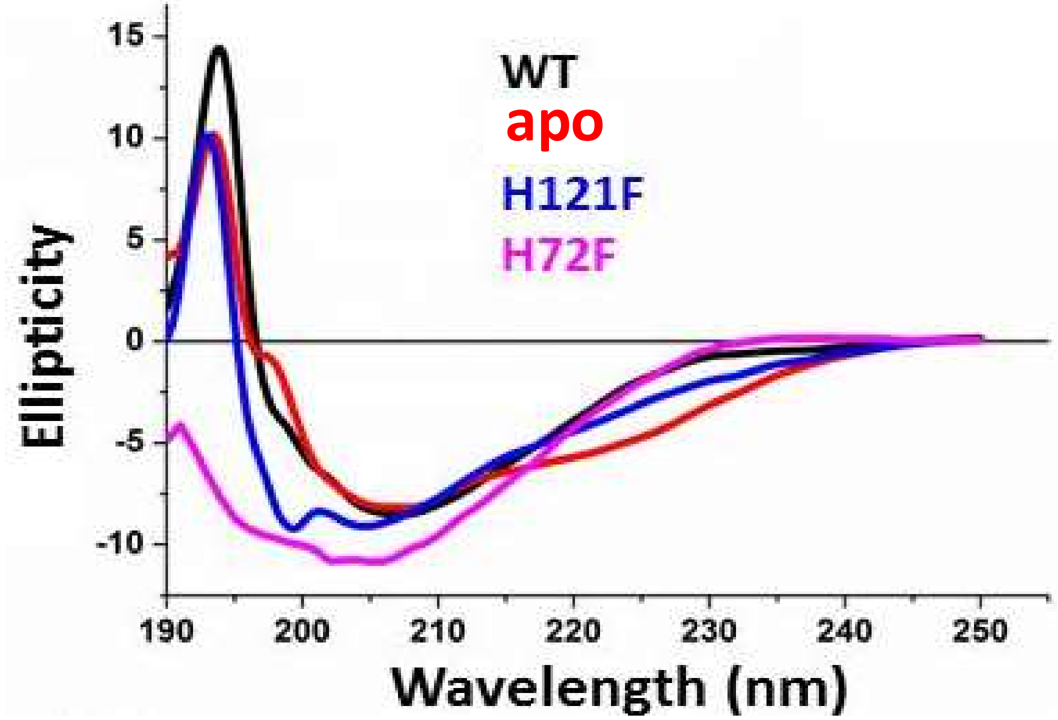


**Figure S3** Far UV-CD spectra of the SOD1 variants.

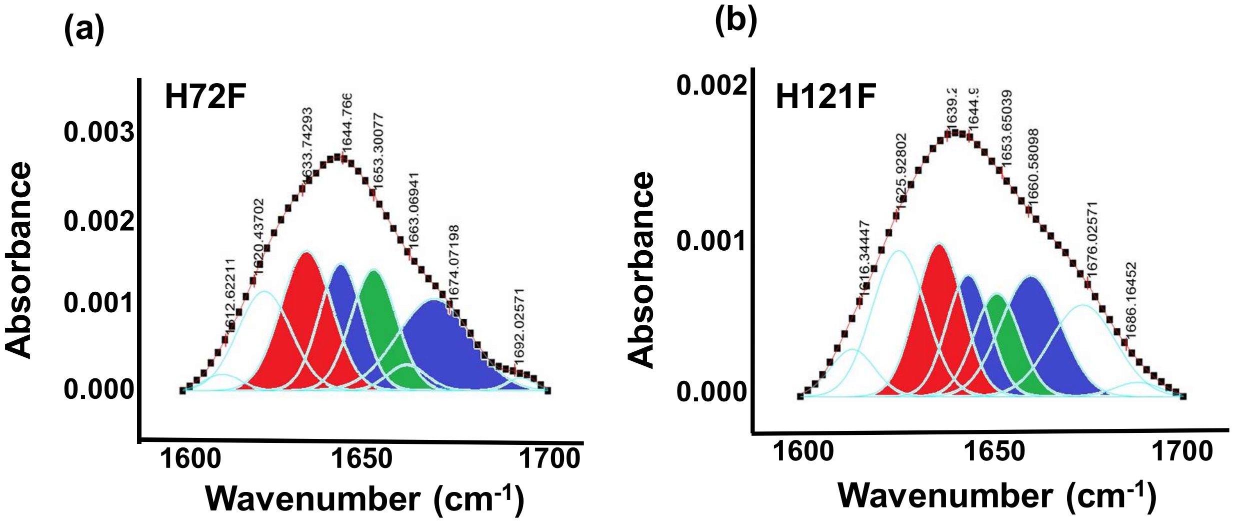


**Figure S4** Deconvoluted FTIR spectra of the amide I region of C=O bond vibrations in case of (a) H72F and (b) H121F. Red contour(~ 1637 $cm^{-1}$ ) indicates beta sheet,blue color contour stands for disorder(1644 $cm^{-1}$) and loops and turns(~1667 $cm^{-1}$); green contour represents alpha helical character(~1653$cm^{-1}$).

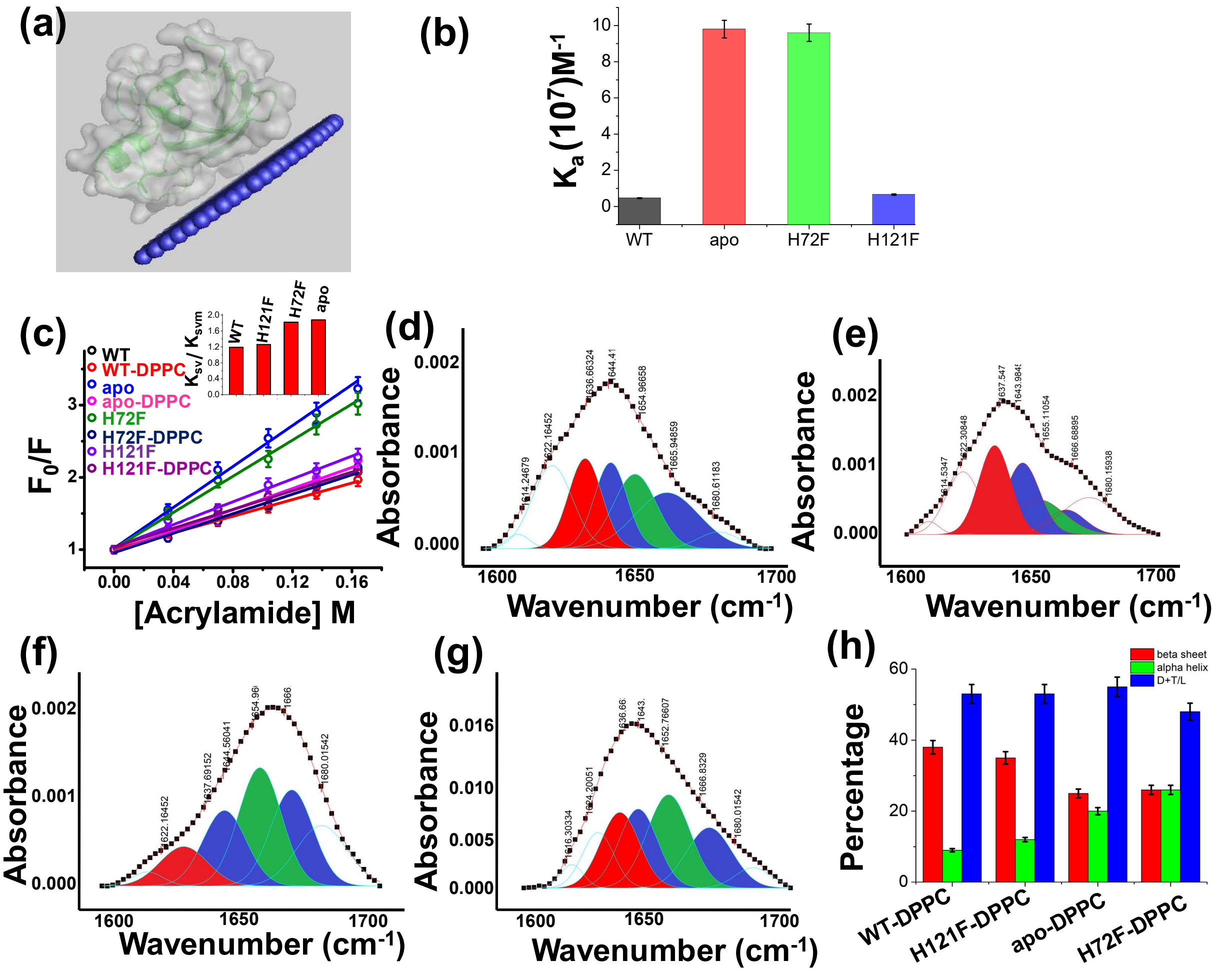

**Figure S5** (a) Membrane attachment of WT SOD1 with membrane as predicted with OPM. (b) Binding constants ($K_a$) of WT,apo, H121F and H72F with DPPC SUVs as obtained from FCS experiments. (c) Acrylamide quenching experiments to probe the solvent exposure of Tryp 32 in the presence and absence of membranes. The inset shows the ratio of $K_{sv}$ and $K_{svm}$ for the four SOD1 variants. Deconvoluted FTIR spectral signatures of (d) WT,(e) H121F,(f) apo and (g) H72F in membrane bound conditions. The red region indicates the beta sheet conformation, blue color coded region stands for the disorder and loops and turns conformations and the green color coded region denotes the alpha helical content.(h) Extent of secondary structures of the SOD1 variants in the presence of membranes as evaluated from FTIR spectroscopy.

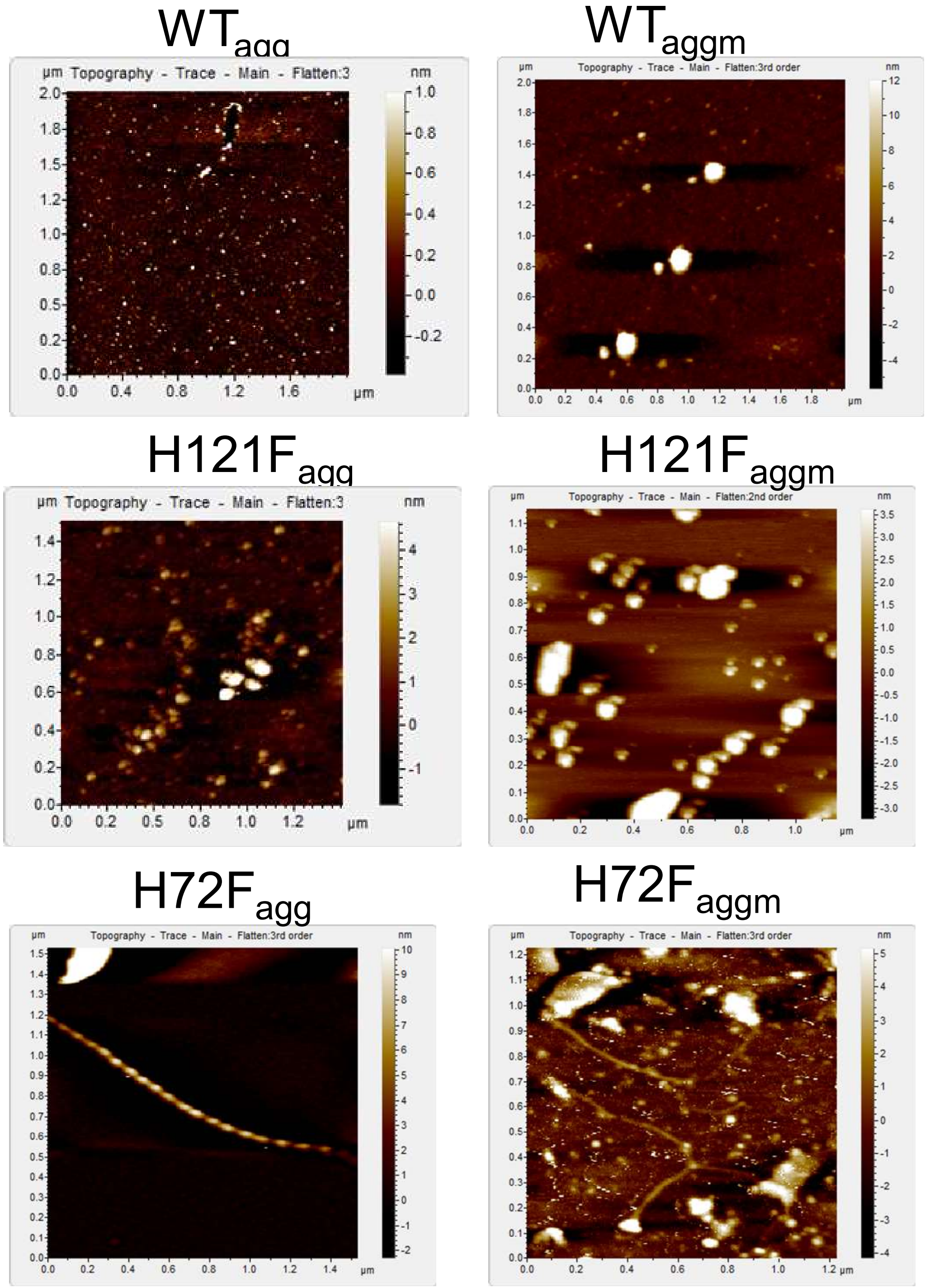


**Figure S6** AFM topographic images of the aggregates of different protein samples at the final points of aggregation.

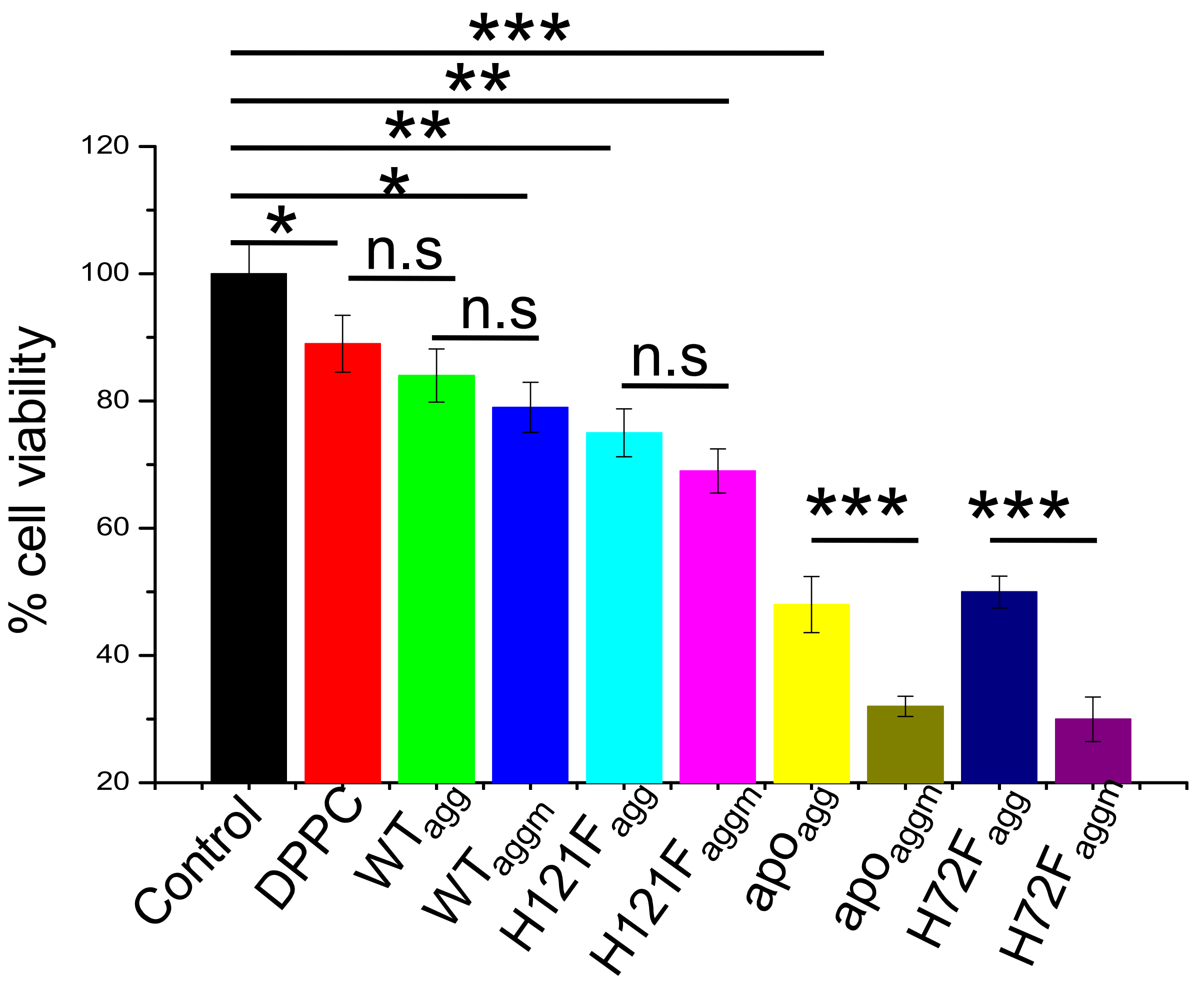


**Figure S7** MTT assay to detect the cell viability of neuronal cells (SHSY5Y) when treated with different aggregates of SOD1 protein variants. The concentration of the aggregate for each sample was taken 5 μM. Error bar indicates the standard deviation. n.s stands for nonsignificant data. For the significant changes-- * , P value<0.05; **, P value<0.01; ***, P value< 0.001.

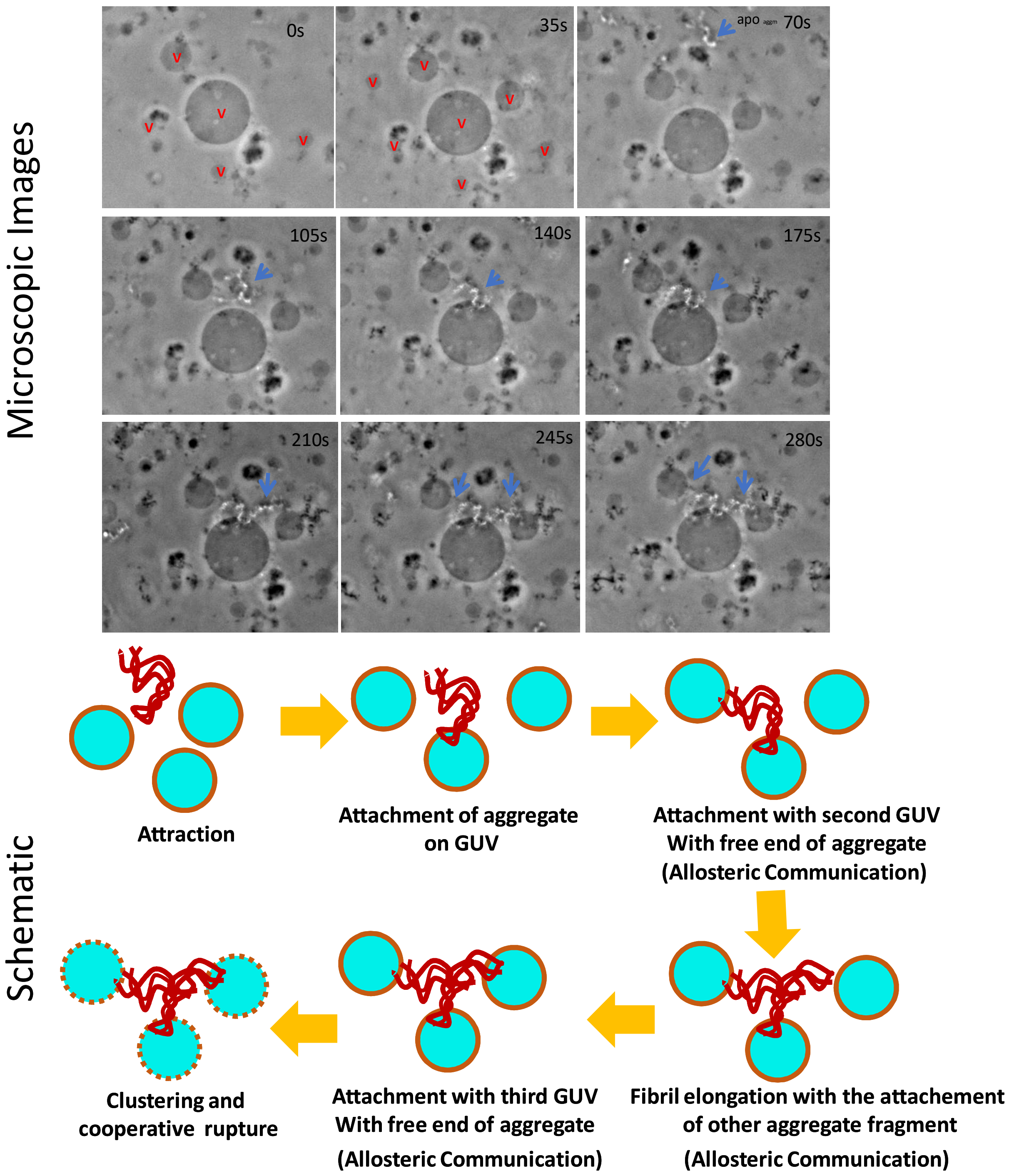


**Figure S8** Time scale optical microscopic images of GUVs when these were treated with apo $_{aggm}$. The size of the central vesicle is ~30 µm. Stepwise deformation kinetics and GUVs-clustering by aggregates were shown in the image panel as well as in the schematic diagram. The observation suggested that the attachment of a fibril occurred first on a GUV surface. Then the free site of the aggregate interacted with other GUV of the proximity presumably through allosteric communication mechanistic way. Next, other small fibrillar aggregate species got attached with the previous GUV attached fibril for elongation and there after it got bound with nearest another vesicle. Finally, vesicle clustering happened by the aggregates and GUV deformation/ rupture took place. Here ‘v’ stands for vesicle.

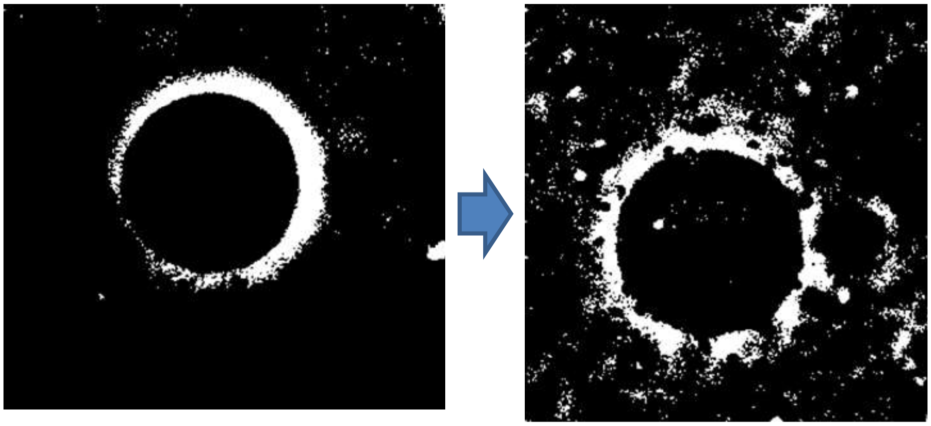

**Figure S9** High contrast images of the GUV when it was treated with $apo_{agg}$. The use of $apo_{agg}$ also resulted in vesicle fusion, but the extents and the number of participating vesicles were found to be more in the case of $apo_{aggm}$ (Fig.4f).

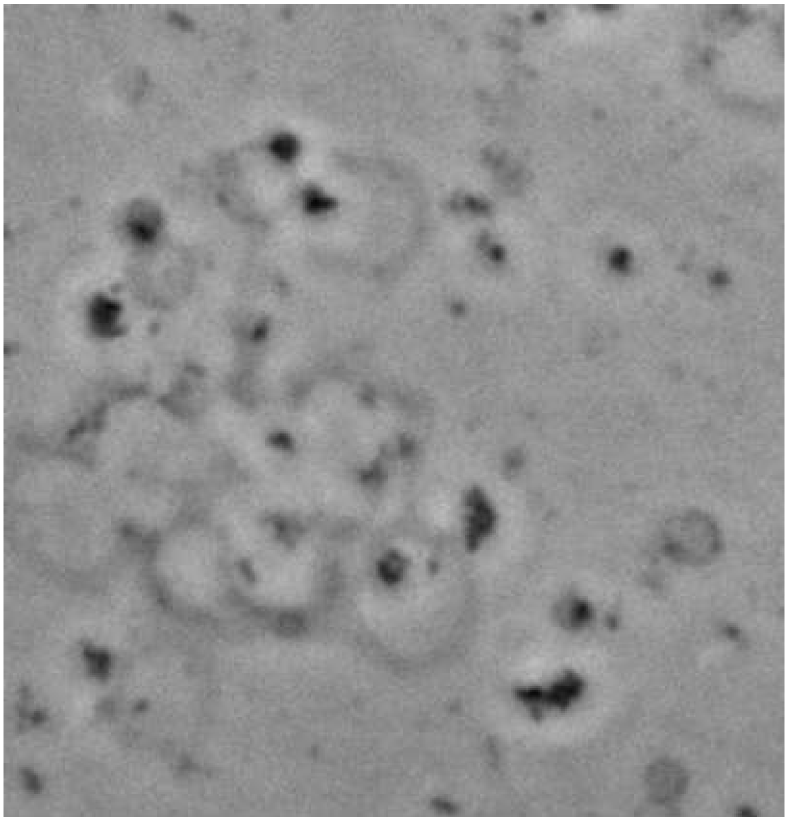

**Figure S10 Assembly of vesicles/ GUV clustering when GUVs were treated with H72F $_{aggm}$.**

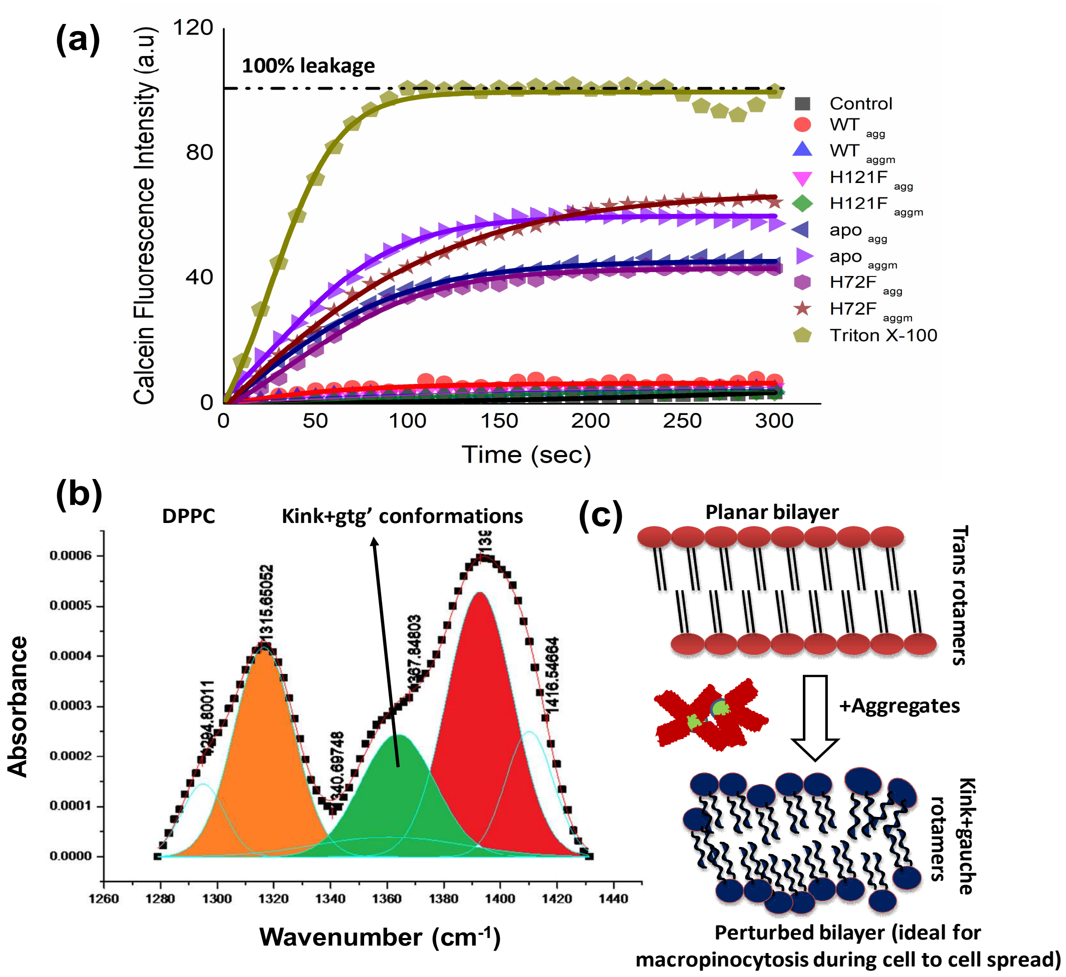


**Figure S11** (a) Calcein release assay to probe the membrane rupture and pore formation in SUVs mimicking the synaptic vesicle composition (DOPC:DOPE:DOPS in the ratio 2:5:3 ) while these SUVs were treated with different protein aggregates. The full release/rupture of the synaptic vesicle mimicking SUVs in presence of Triton X-100 was considered as 100% release with respect to which other aggregate induces leakage percentages were calculated. Here 'control' indicates the kinetics in absence of any additive. The fitting of the kinetics were done by exponential growth kinetics equation. (b) FTIR spectral signatures of the $CH_2$ wagging band frequency of DPPC (1280-1440 $cm^{-1}$). The green region (1637$cm^{-1}$) stands for the population of nonplanar gauche and kink conformers which were found to be got increased after the interaction of trans planar bilayer with $apo_{agg}$/ $apo_{aggm}$and $H72F_{agg}$/ $H72F_{aggm}$. It is to be noted that $apo_{aggm}$ and $H72F_{aggm}$ show higher population of nonplanar rotamers than $apo_{agg}$ and $H72F_{agg}$ respectively. On the other hand, insignificant changes were found when trans-planar bilayer was treated with $WT_{agg}$, $WT_{aggm}$ and $H121F_{agg}$ , $H121F_{aggm}$. (c) Schematic representation of the planar membrane perturbations by aggregates and the changes from trans to gauche rotamers of the hydrocarbon tail region. This kind of lipid's hydrocarbon conformational change may induce the macropinocytosis in membrane by aggregate induced increased flexibility in the membrane.

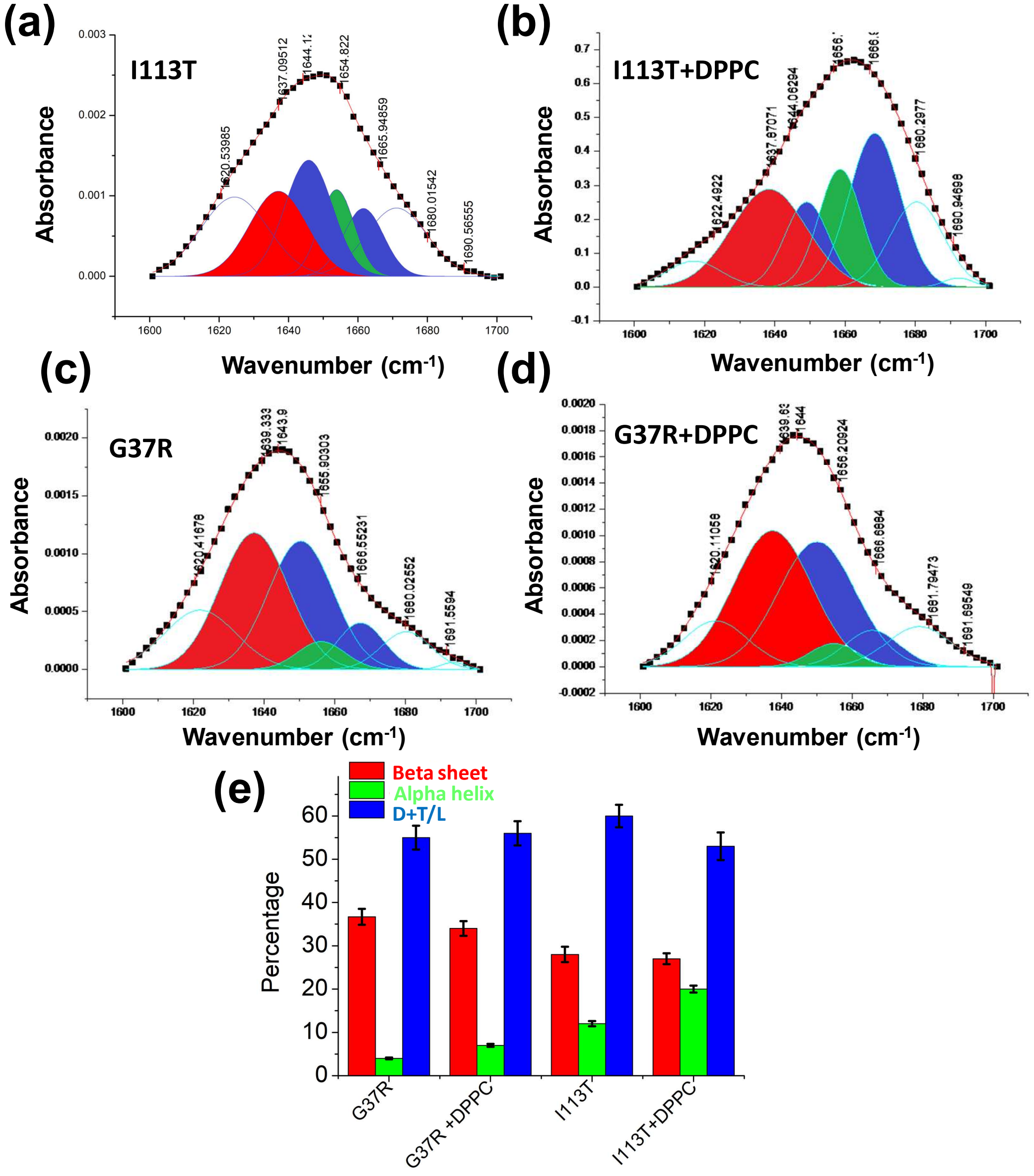


**Figure S12** (a)-(d) Deconvoluted FTIR spectral signatures which correspond to the carbonyl vibrational frequency of amide-I region (1600-1700 $cm^{-1}$) of I113T and G37R mutant both in absence and presence of lipid membranes. (e) perecentage of secondary conformations of G37R and I113T disease mutants both in absence and presence of DPPC SUVs.

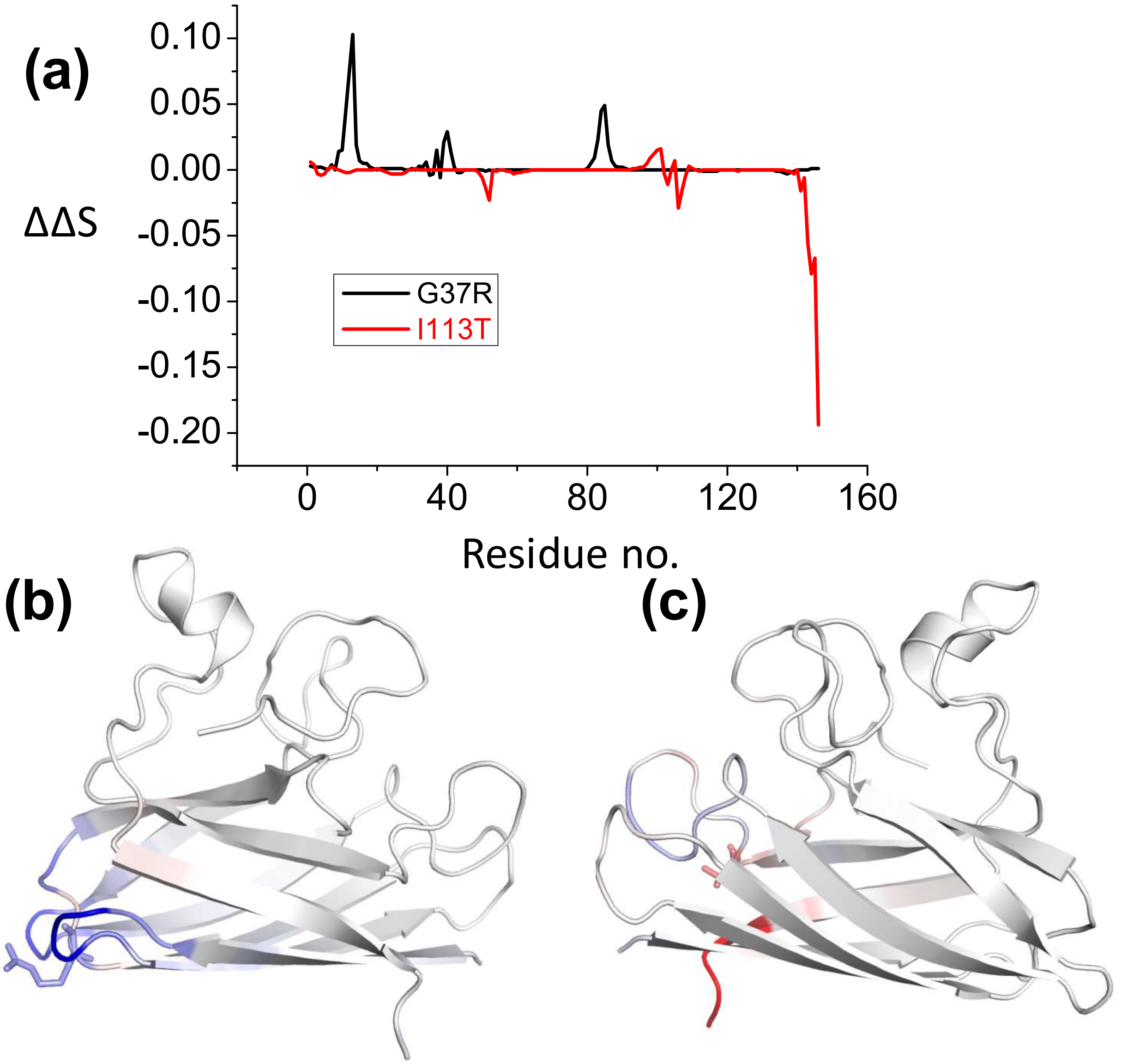


**Figure S13** Effect of point mutations (here disease mutations) of the proteins flexibility, conformational stability and dynamics as predicted using DynaMut webserver (http://biosig.unimelb.edu.au/dynamut/). (a) residue wise vibrational entropy changes for G37R and I113T; (b) the structure of G37R indicating the increase in the structural rigidity (blue color indicates the region becomes rigid due to the mutation); (c) structural flexibility increases in I113T mutant (red color indicates the region becomes more flexible due to mutation).

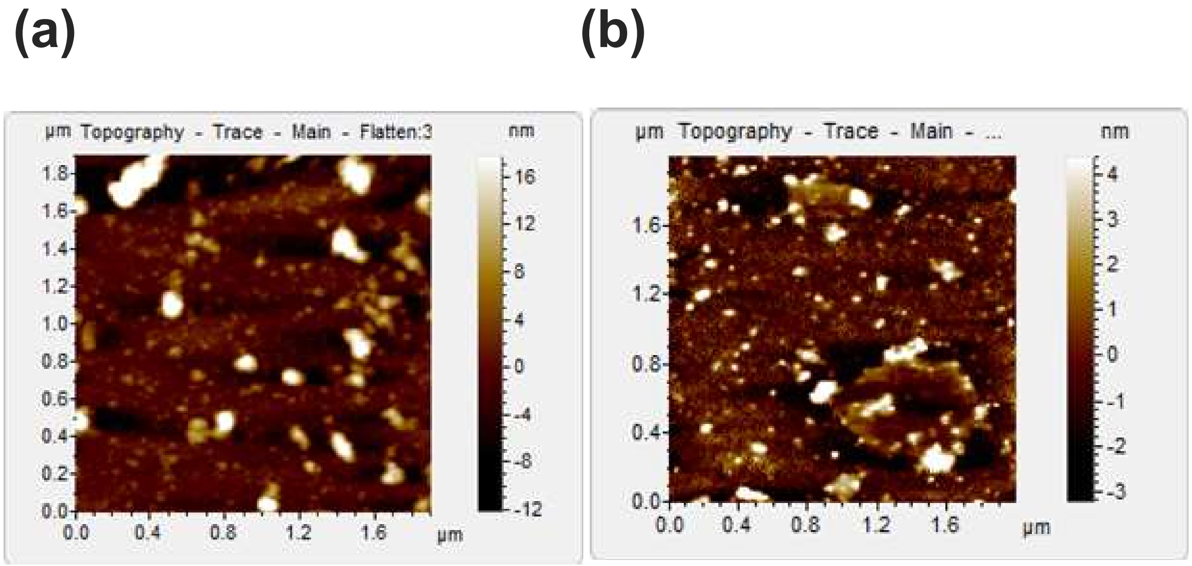


**Figure S14** AFM topographic images of the aggregates of G37R that was formed in absence (a) and presence (b) of membrane.

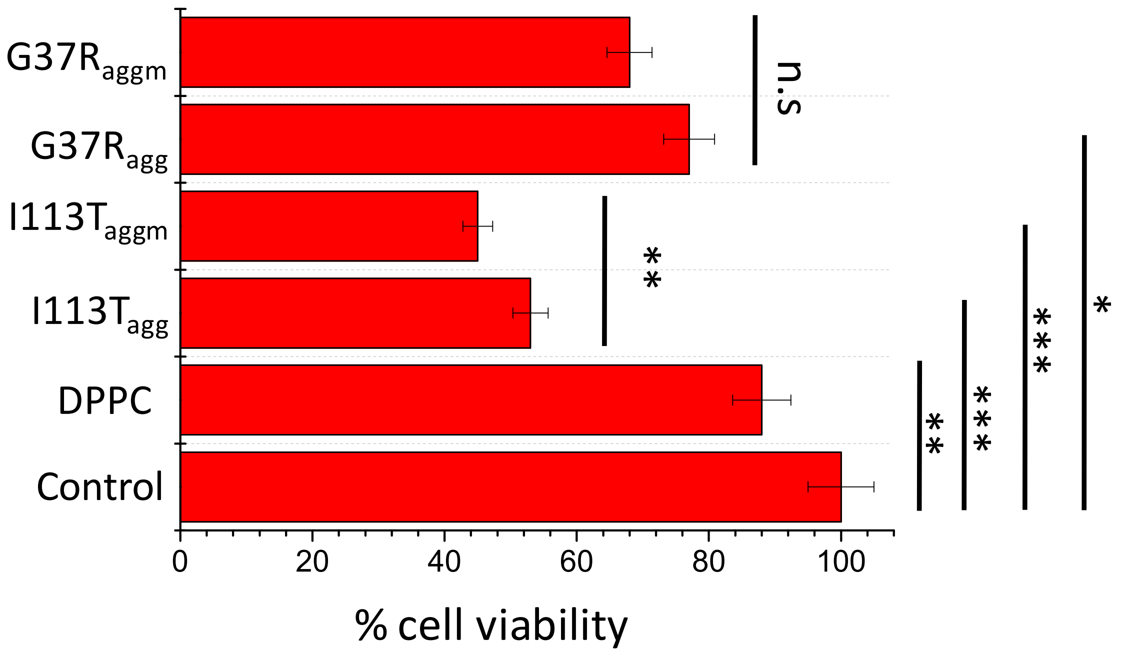


**Figure S15** MTT assay to detect the cell viability of neuronal cells (SHSY5Y) when treated with different aggregates of SOD1 disease mutants. The concentration of the aggregate for each sample was taken 5 μM. n.s stands for nonsignificant data. For the significant changes—,'*' P value<0.05; **, P value<0.01; ***, P value< 0.001. Error bar indicates the standard deviation for the triplicate experiments.

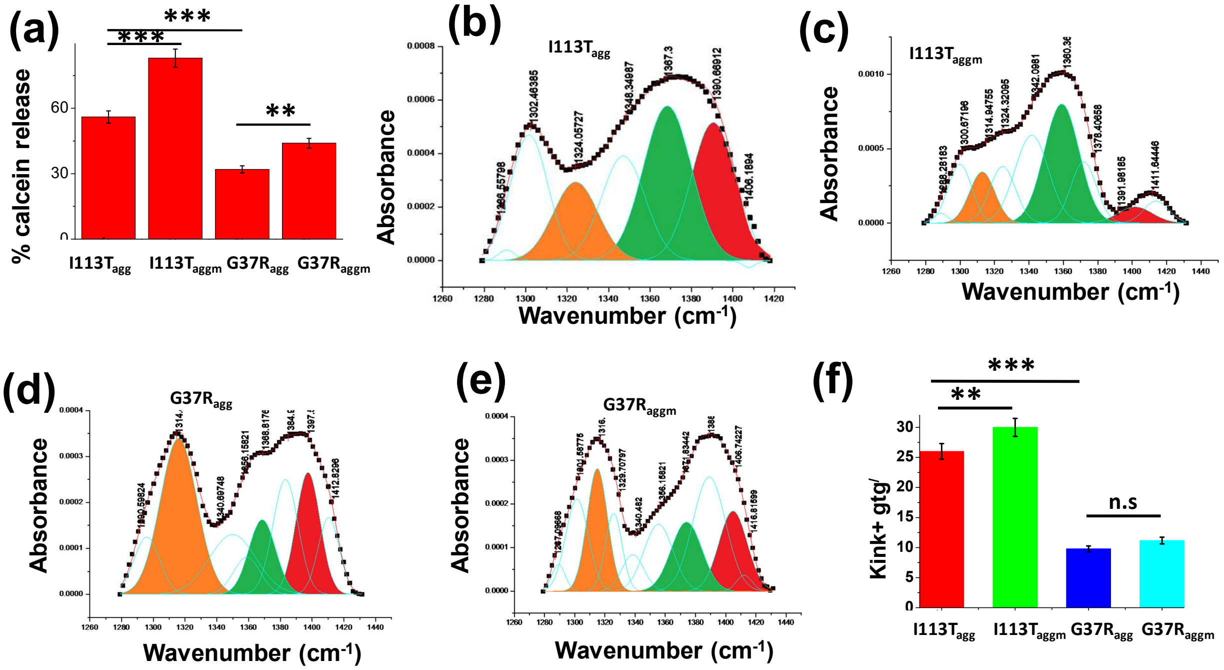


**Figure S16** (a) Calcein release percentage when dye entrapped Small Unilamellar Vesicles (SUVs) were treated with the aggregates of I113T and G37R which were formed in absence and presence of membranes.(b)-(e) show the gaussian fittings of the $CH_2$ wagging band region of planar DPPC membrane after the treatments of different disease mutant aggregates. (f) Populations of nonplanar conformations in membrane bilayer which is perturbed by the aggregates of I113T and G37R which are both formed in absence and presence of membranes. Here n.s denotes nonsignificant whereas ** is significant with p values<0.01 and *** is highly significant with p values<0.001. $I113T_{agg}$ and $I113T_{aggm}$ indicate the aggregates of I113T protein which are formed in absence and presence of DPPC SUVs. $G37R_{agg}$ and $G37R_{aggm}$ indicate the aggregates of G37R protein which are formed in absence and presence of DPPC SUVs. Error bar indicates the standard deviations of the values obtained from experiments those were performed triplicate.

### Supporting References:


1. Wright GS, Antonyuk SV, Kershaw NM, Strange RW, Hasnain SS. Ligand binding and aggregation of pathogenic SOD1. *Nature communications* **4**, 1758 (2013).

2. Marklund S, Marklund G. Involvement of the superoxide anion radical in the autoxidation of pyrogallol and a convenient assay for superoxide dismutase. *The FEBS Journal* **47**, 469-474 (1974).

3. McCord JM, Fridovich I. Superoxide dismutase an enzymic function for erythrocuprein (hemocuprein). *Journal of Biological chemistry* **244**, 6049-6055 (1969).

4. Lakowicz J. Principles of fluorescence microscopy.). Kluwer Academic, New York (1999).

5. Bandekar J. Amide modes and protein conformation. *Biochimica et Biophysica Acta (BBA)-Protein Structure and Molecular Enzymology* **1120**, 123-143 (1992).

6. Stewart JCM. Colorimetric determination of phospholipids with ammonium ferrothiocyanate. *Analytical biochemistry* **104**, 10-14 (1980).

7. Benachir T, Monette M, Grenier J, Lafleur M. Melittin-induced leakage from phosphatidylcholine vesicles is modulated by cholesterol: a property used for membrane targeting. *European biophysics journal* **25**, 201-210 (1997).

8. Pott T, Bouvrais H, Méléard P. Giant unilamellar vesicle formation under physiologically relevant conditions. *Chemistry and physics of lipids* **154**, 115-119 (2008).

9. Nandi R*, et al.* A novel nanohybrid for cancer theranostics: folate sensitized Fe 2 O 3 nanoparticles for colorectal cancer diagnosis and photodynamic therapy. *Journal of Materials Chemistry B* **5**, 3927-3939 (2017).

10. Kundu A, Kundu S, Chattopadhyay K. The presence of non-native helical structure in the unfolding of a beta-sheet protein MPT63. *Protein Science* **26**, 536-549 (2017).

11. Chattopadhyay K, Saffarian S, Elson EL, Frieden C. Measurement of microsecond dynamic motion in the intestinal fatty acid binding protein by using fluorescence correlation spectroscopy. *Proceedings of the National Academy of Sciences* **99**, 14171-14176 (2002).

12. Wako H, Saitô N. Statistical mechanical theory of the protein conformation. II. Folding pathway for protein. *Journal of the Physical Society of Japan* **44**, 1939-1945 (1978).

13. Muñoz V, Eaton WA. A simple model for calculating the kinetics of protein folding from three-dimensional structures. *Proceedings of the National Academy of Sciences* **96**, 11311-11316 (1999).

14. Gopi S, Aranganathan A, Naganathan AN. Thermodynamics and folding landscapes of large proteins from a statistical mechanical model. *Current Research in Structural Biology* **1**, 6-12 (2019).

15. Naganathan AN. Predictions from an Ising-like statistical mechanical model on the dynamic and thermodynamic effects of protein surface electrostatics. *Journal of chemical theory and computation* **8**, 4646-4656 (2012).

16. Rajasekaran N, Gopi S, Narayan A, Naganathan AN. Quantifying protein disorder through measures of excess conformational entropy. *The Journal of Physical Chemistry B* **120**, 4341-4350 (2016).

17. Sekhar A*, et al.* Thermal fluctuations of immature SOD1 lead to separate folding and misfolding pathways. *Elife* **4**, e07296 (2015).

18. Yang J, Zhang Y. I-TASSER server: new development for protein structure and function predictions. *Nucleic acids research* **43**, W174-W181 (2015).

19. Lomize MA, Lomize AL, Pogozheva ID, Mosberg HI. OPM: orientations of proteins in membranes database. *Bioinformatics* **22**, 623-625 (2006).

20. Lomize AL, Pogozheva ID, Lomize MA, Mosberg HI. The role of hydrophobic interactions in positioning of peripheral proteins in membranes. *BMC Structural Biology* **7**, 44 (2007).
